# Network-based integrative multi-omics approach reveals biosignatures specific to COVID-19 disease phases

**DOI:** 10.1101/2023.09.29.560110

**Authors:** Francis E. Agamah, Thomas H.A. Ederveen, Michelle Skelton, Darren P. Martin, Emile R. Chimusa, Peter A.C. ’t Hoen

## Abstract

**Background:** COVID-19 disease is characterized by a spectrum of disease phases (mild, moderate, and severe). Each disease phase is marked by changes in omics profiles with corresponding changes in the expression of features (biosignatures). However, integrative analysis of multiple omics data from different experiments across studies to investigate biosignatures at various disease phases is limited. Exploring an integrative multi-omics profile analysis through a network approach could be used to determine biosignatures associated with specific disease phases and enable the examination of the relationships between the biosignatures.

**Aim:** To identify and characterize biosignatures underlying various COVID-19 disease phases in an integrative multi-omics data analysis.

**Method:** We leveraged the correlation network approach to integrate transcriptomics, metabolomics, proteomics, and lipidomics data. The World Health Organization (WHO) Ordinal Scale (WOS) was used as a disease severity reference to harmonize COVID-19 patient metadata across two studies with independent data. A unified COVID-19 knowledge graph was constructed by assembling a disease-specific interactome from the literature and databases. Disease-state omics-specific graphs were constructed by integrating multi-omics data with the unified COVID-19 knowledge graph. We expanded on the network layers of multiXrank, a random walk with restart on multilayer network algorithm, to explore disease state omics-specific graphs and perform enrichment analysis.

**Results:** Network analysis revealed the biosignatures involved in inducing chemokines and inflammatory responses as hubs in the severe and moderate disease phases. We observed more shared biosignatures between severe and moderate disease phases as compared to mild-moderate and mild-severe disease phases. We further identified both biosignatures that discriminate between the disease states and interactions between biosignatures that are either common between or associated with COVID-19 disease phases. Interestingly, cross-layer interactions between different omics profiles increased with disease severity.

**Conclusion:** This study identified both biosignatures of different omics types enriched in disease-related pathways and their associated interactions that are either common between or unique to mild, moderate, and severe COVID-19. These biosignatures include molecular features that underlie the observed clinical heterogeneity of COVID-19 and emphasize the need for disease-phase-specific treatment strategies. In addition, the approach implemented here can be used for other diseases.

**Key findings:**

⍰ Integrative multi-omics analysis revealed biosignatures and biosignature interactions associated with COVID-19 disease states.
⍰ Disease severity increases with biosignature interactions across different multi-omics data.
⍰ The harmonization approach proposed and implemented here can be applied to other diseases

## Background

Coronavirus Disease-2019 (COVID-19) is a contagious respiratory disorder caused by Severe Acute Respiratory Syndrome Coronavirus 2 (SARS-CoV-2), a newly emerged β coronavirus belonging to the Coronaviridae family (1). Since its discovery in Wuhan, China, in December 2019, COVID-19 established itself as a devastating global pandemic that has created disruptions across healthcare, economic, and social systems (2).

COVID-19 is characterized by a range of clinical phenotypes that reflect the spectrum of disease severity (i.e., mild, moderate, and severe herein defined as disease phases). Disease phenotypes are broadly classifiable as asymptomatic and symptomatic with approximately 85% of infected patients (vaccinated and non-vaccinated) showing mild to moderate symptoms and approximately 15% of infected patients suffering from potentially life-threatening complications (3). The mild to moderate disease phase includes disease conditions with few or no infection symptoms, hospitalization with either no oxygen therapy required or with oxygen given by mask or nasal prongs, and no fatalities. In contrast, the severe disease phase includes disease conditions that, besides death, could involve one or a combination of hospitalization, oxygen therapy involving mechanical ventilation, respiratory failure, and significant immune dysregulation (1).

Each COVID-19 disease phase (i.e., mild, moderate, and severe) is marked by changes in omics profiles with corresponding changes in the expression levels of biosignatures (4–6). In the context of this research, we define biosignatures as omics features that include proteins, transcripts, lipids, and metabolites. Although some of these biosignatures have a connection to the pathology of COVID-19 illness, not all of them actively impact the expressed disease phenotype when they are dysregulated. Different major dysregulated biosignatures are linked to host responses to COVID-19 (7–10). Such biosignatures may serve not only as potential biomarkers to stratify patients according to disease severity and/or provide detailed prognostic information but could also contribute to the development of treatments that are more specifically targeted at particular disease states.

Several individual-omics (11–21) and multi-omics COVID-19 investigations (7, 22-28) have identified biosignatures that are associated with disease progression. Individual omics studies provide specific insights into the contributions/manifestations of biosignatures at that omics level during disease progression but have not accounted for the impact(s) of other omics layers. Multi-omics studies present a means to collectively compare multiple omics data from different experiments either on the same samples or across studies, to yield a more holistic understanding of the biochemical underpinnings of COVID-19 outcomes.

However, few of these omics studies (7, 22-28) focusing on identifying the biochemical drivers of COVID-19 clinical heterogeneity have computationally integrated multi-omics data from different study samples with existing biological knowledgebases to explore biosignatures of different omics types and their connections across different disease phases. We suggest that this kind of multi-omics data integration is essential when attempting to explain the molecular dynamics underpinning the heterogeneity of COVID-19 infections while accounting for both prior knowledgebase and data from independent studies.

Network-based integrative approaches have revolutionized multi-omics analyses by providing the framework to build on existing knowledgebases when using new data to infer interactions between multiple different omics profiles within the context of a graph representation (29). This has been shown to provide the opportunity not only to elucidate interactions that can occur among all classes of biomolecules in a biological system but also, to prioritize biosignatures that could discriminate disease severity. The approach represents the biomolecules that are most indicative of differences between disease states (i.e. the biosignatures) as nodes in the graph and infers relationships between them. For this reason, we hypothesized that (i) investigating biosignatures across different phases of COVID-19 disease will provide insights into the molecular underpinnings of the enormous clinical heterogeneity of COVID-19 and (ii) associations between biosignatures within a biologically meaningful network would permit the prioritization of biosignatures that discriminate between the disease states and could yield leads for potential drug targets.

Our study implemented a multi-layered network-based approach to identify and characterize biosignatures underlying various COVID-19 disease phases by integrating transcriptomics, metabolomics, proteomics, and lipidomics data (each representing a different data layer) with known interactome data (i.e. our present knowledgebase). We build disease-state omics-specific graphs and apply a network diffusion-based method to predict biosignatures and their associated interactions - both within and between data layers - that are linked to the various COVID-19 disease phases.

The major contributions of this work are: (1) a new method of harmonizing patient disease severity metrics by leveraging the World Health Organization (WHO) Ordinal Scale (WOS)and patient metadata; (2) the assembly of a unified COVID-19 knowledge graph from different curated sources of interactome data (our resent knowledgebase); (3) the construction of a disease-state omics-specific graph by integrating curated transcriptomics, proteomics, lipidomics, and metabolomics datasets from two different multi-omics studies; and (4) identified biosignatures and their associated interactions that are shared and/or unique to mild, moderate, and severe COVID-19 disease-states. Our study identifies biosignatures that discriminate between disease states and also shows that COVID-19 disease severity increases together with the numbers of inferred interactions across different omics layers

## Results

### Harmonized clinical severity between patients’ metadata

The clinical severity harmonization was a crucial step in integrating and analyzing data from multiple sources and platforms to gain a more comprehensive understanding of biological systems and disease mechanisms. There were (**Figure 2**) more severe samples than any other in the Overmyer et al. (25) dataset followed by mild and then moderate samples. For the Su et al., (24) dataset, there were similar numbers of mild and moderate samples and fewer severe samples. The classified samples (**Supplementary data Table 2**) were used to split the omics counts data into disease states before constructing the disease-state omics-specific graphs.

**Figure 1.**
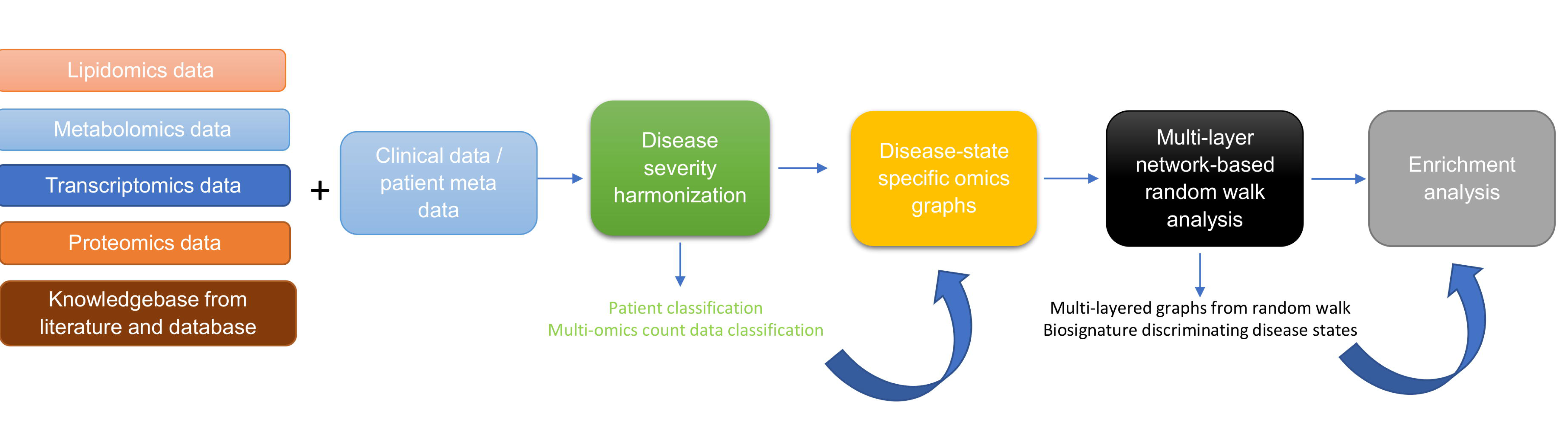
Diagram illustrating the workflow implemented in this study. The workflow begins with curating lipidomics, metabolomics, transcriptomics, proteomics data and their associated patient metadata and knowledge graph from literature and databases. Next, we leveraged on the patient metadata to perform disease severity harmonization. To harmonize the clinical severity of patients, we used the WOS as the reference for classifying disease severity into three disease states, such that: (1) mild disease state represents COVID-19 patients with WOS 1-2, (2) moderate disease state represents COVID-19 patients with WOS 3-4, and (3) severe disease state represents COVID-19 patients with WOS 5-9. We then used the harmonized information to split the omics datasets according to disease severity prior to constructing disease-state omics-specific graphs. We then performed randomwalk analysis on the graphs to predict biosignatures discriminating the various disease states. Finally we performed enrichment analysis.

**Figure 2.**
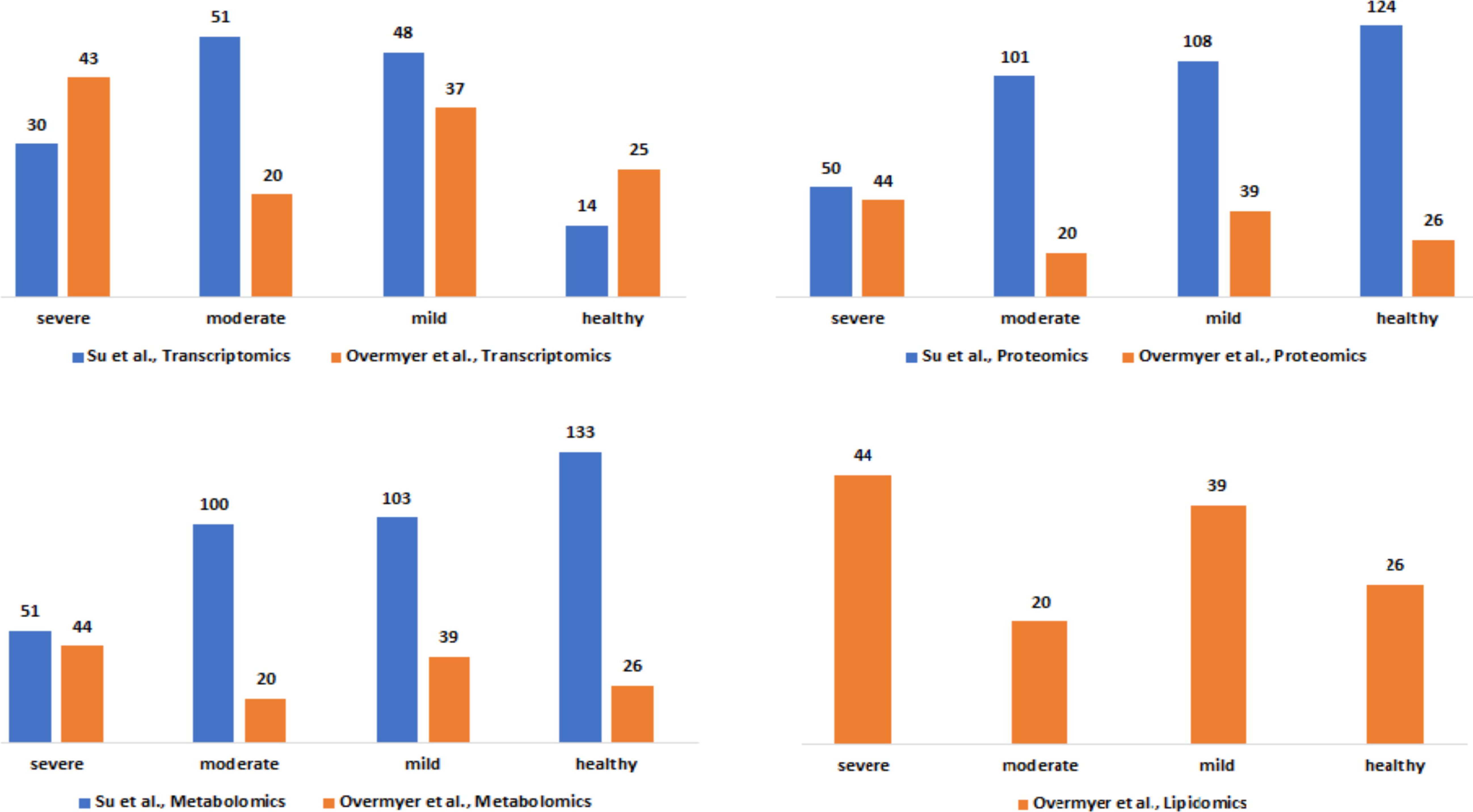
Description of the Su et al., and Overmyer et al., samples based on the omics data type and the disease severity levels after the harmonization process.

### Integrative network-based multi-omics analysis

#### Construction of disease-state omics-specific graphs

We constructed a unified knowledge graph (**Figure 3A**) comprising four different edge types by merging the protein-protein interaction data from GeneMANIA, metabolite-metabolite interactome, lipid-lipid interactome, and the extracted data from the COVID-19 knowledge graph. Importantly, the unified knowledge graph formed the basis for integrating the processed multi-omics data (**Figure 3B**) and constructing disease-state omics-specific graphs for mild, moderate, and severe COVID-19 disease states (**Supplementary File 1**). Additionally, the disease-state omics-specific graphs enabled us to investigate COVID-19 disease states in the context of specific omics data types.

**Figure 3.**
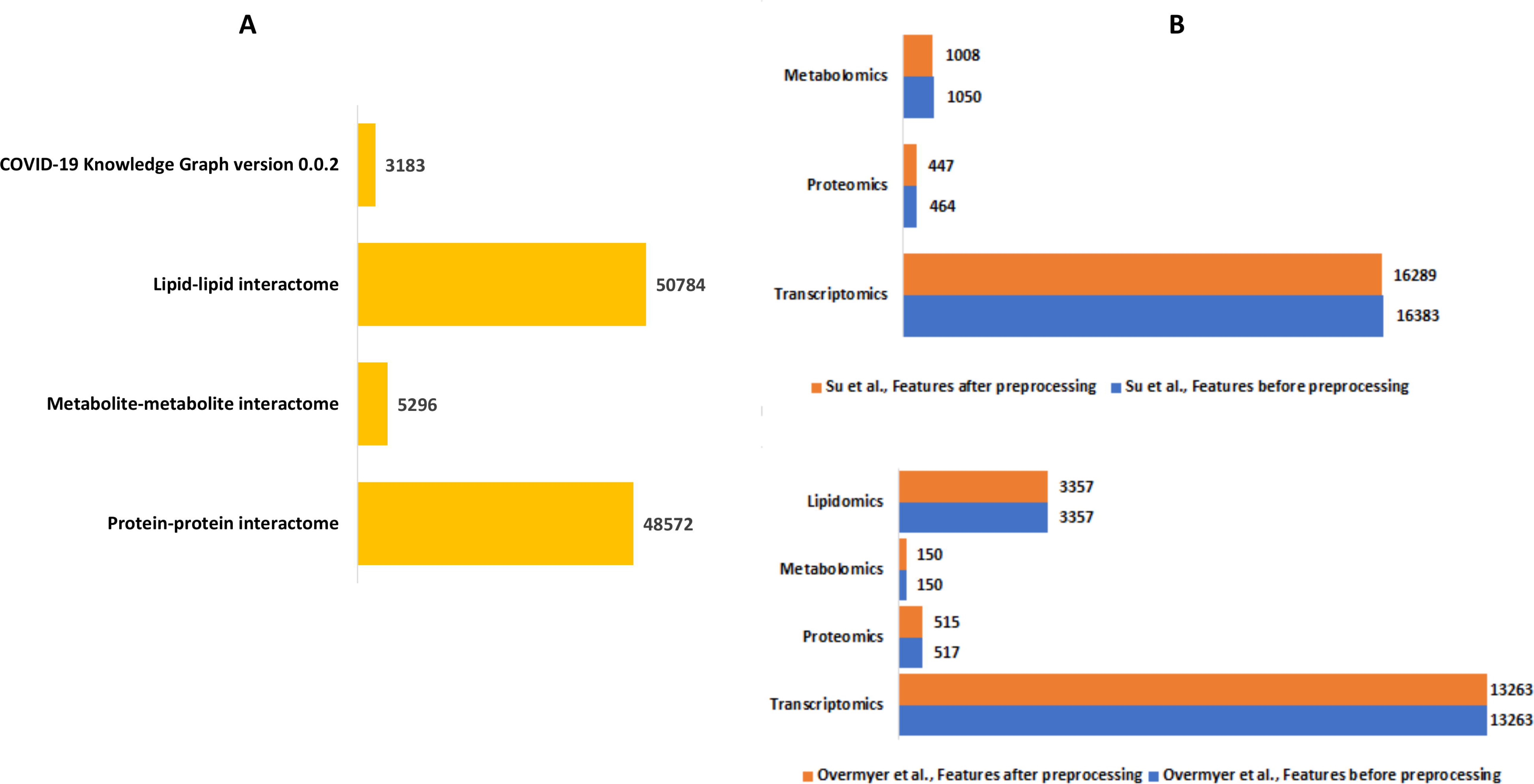
**(A)** Summary of the edge count in the interactome datasets used to construct the unified knowledge graph **(B)** Summary of the number of features in the omics experimental datasets before and after data processing.

#### Identified seed nodes for network exploration

The random walk method is a technique for detecting the spread of biological information throughout networks. The concept behind the random walk method is such that a hypothetical particle exploring the network structure starts from a seed node (29). We selected the seed node at which random walks began (**Table 1**) using both data-driven and hypothesis-driven approaches. Whereas for the data-driven approach, we selected a seed based on a computed integrated node centrality metric score (**Supplementary data**), for the hypothesis-driven approach, we selected nodes based on previously reported associations of molecular features with different COVID-19 disease states.

**Table 1.** Selected seeds for random walk network exploration.

| Approach | Seed node | Integrated centrality score | Feature type |
| --- | --- | --- | --- |
| Data-driven | STAT1 | 53529.0403 | Transcript |
|  | SOD2 | 2215.5746 | Protein |
|  | 3-hydroxyoctanoate | 1506.9998 | Metabolite |
|  | Unknown_mz_815.61548+_RT_27.063 | 9936.9781 | Lipid |
| Hypothesis-driven | IL6R | 252.0804 | Transcript |
|  | IL6 | 2208.0288 | Protein |

**Table 2.**
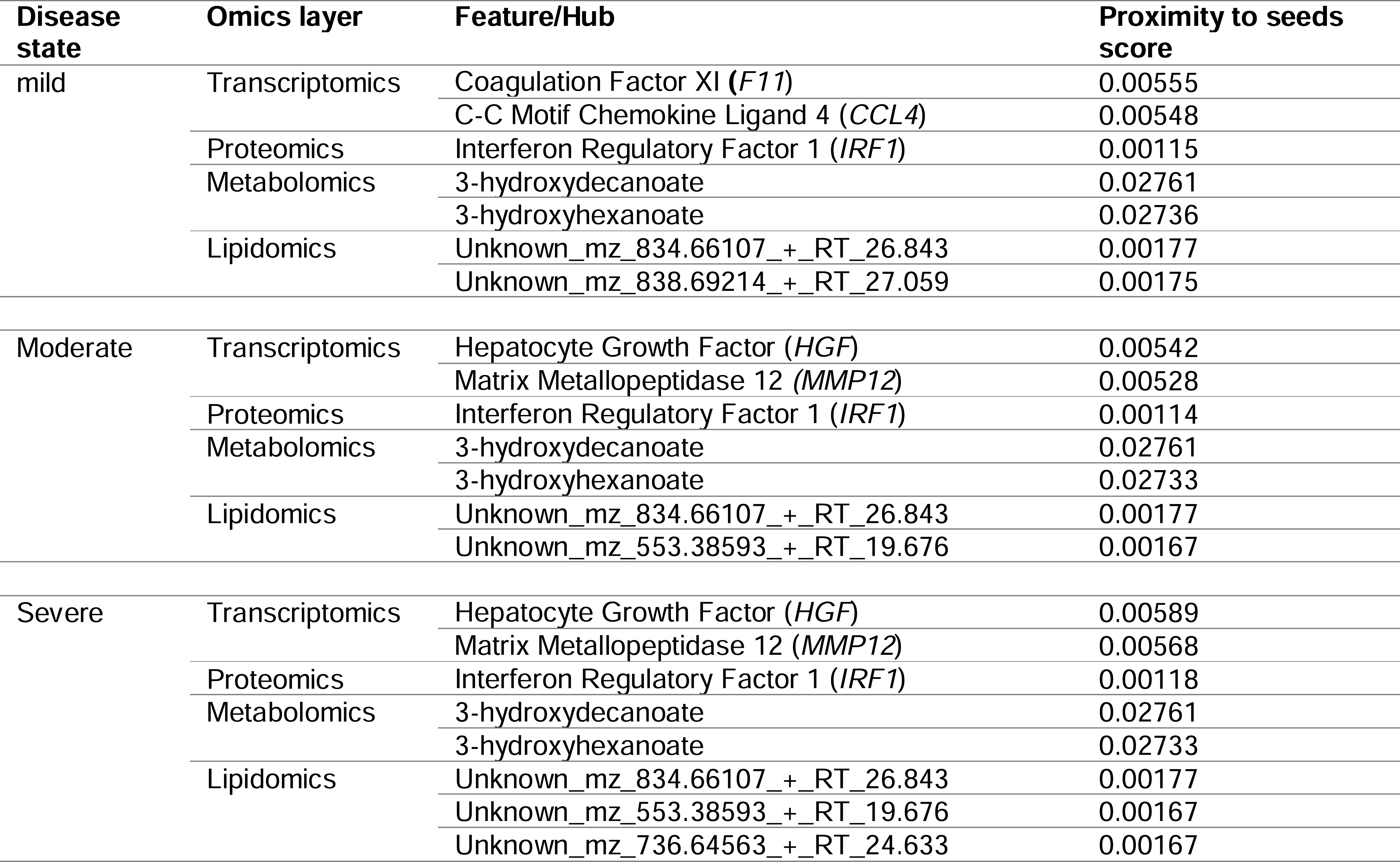
Key hubs identified in the disease-state omics-specific graph upon using seeds from the data-driven approach.

For the data-driven approach, one selected seed node was the Signal Transducer And Activator Of Transcription 1 (*STAT1*) from the transcriptomics layer. *STAT1* is known to be involved in immune responses and antiviral activity (30) and is reported to be upregulated in mild and severe COVID-19 cases, with the phosphorylation of the gene highly enhanced in severe disease states (31). Another selected seed node from the proteomics layer was Superoxide Dismutase 2 (*SOD2*), an essential antioxidant enzyme that protects cells from superoxide radical anions which are known to be significantly under-expressed in the plasma (32), and in the lung cells of severe COVID-19 patients (33). From the metabolomics layer, we used 3-hydroxyoctanoate, as a seed: 3-hydroxyoctanoate is a metabolite of medium-chain fatty acid oxidation that has been identified as a marker for primary defects of beta-hydroxy fatty acid metabolism and it is a conjugate acid to 3-hydroxyoctanoate, a biomarker of asymptomatic COVID-19 infection that is involved in important pathways such as the activation of macrophage and platelet aggregation [33]. From the lipidomics layer, we identified Unknown_mz_815.61548_+_RT_27.063, an uncharacterized lipid as a seed.

For the hypothesis-driven approach, we selected *interleukin-6 (IL-6)* and *IL-6R* as seeds. Besides the pathologic roles of these molecules in immune-inflammatory diseases such as COVID-19, it has been hypothesized that inhibition of *IL-6* receptors (*IL-6Rs*) by tocilizumab ameliorates the symptoms of severe COVID-19 and reduces mortality (34, 35). We aimed to find out how *IL-6* and *IL-6R* influence disease severity.

#### Random walk analysis on disease-state omics-specific graphs using data-driven seeds

We used various omics features (**Table 1**) as seed nodes for the random walk analysis. The features in each disease-state omics-specific graph were ranked by their proximity to the seeds. The generated multi-layered graphs (accessible at http://cytoscape.h3africa.org), describing the exploration of the seeds during the random walk analysis for each disease state, suggested that cross-layer interactions between the different omics data types influences disease severity. This is evidenced particularly in the protein-metabolite (e.g., *HGF* and 1-palmityl-GPC, *HGF* and 6-bromotryptophan), transcript-metabolite (e.g. *CCL2* and taurine, *CCL2* and 1-(1-enyl-palmitoyl)-2-oleoyl-GPC)), and protein-transcript (e.g. *HGF* and *HLA-B*) interactions. For each omics layer, we identified highly connected features (**Table 3**) forming large subnetworks and defined these as “key hubs”.

**Table 3.** Key hubs identified in the disease-state omics-specific graph upon using seeds from the hypothesis-driven approach.

| Disease state | Omics layer | Feature | Proximity to seeds score |
| --- | --- | --- | --- |
| mild | Transcriptomics | C-C Motif Chemokine Ligand 4 (*CCL4*) | 0.00659 |
|  |  | C-C Motif Chemokine Ligand 2 (*CCL2*) | 0.005629 |
|  | Proteomics | Interleukin 7 Receptor (*IL7R*) | 0.00368 |
|  | Metabolomics | tricosanoyl sphingomyelin (d18:1/23:0)* | 0.0 |
|  |  | suberate (C8-DC) | 0.0 |
|  |  | stearylcholine* | 0.0 |
|  | Lipidomics | Unknown_mz_765.5752_- RT_20.235 | 0.0 |
|  |  | Unknown_mz_765.3349_- RT_6.822 | 0.0 |
|  |  | Unknown_mz_765.3349_+ RT_6.842 | 0.0 |
| Moderate | Transcriptomics | Nuclear Factor Kappa B Subunit 1 (*NFKB1*) | 0.00545 |
|  |  | Interleukin 10 (*IL10*) | 0.00539 |
|  | Proteomics | Interleukin 7 Receptor (*IL7R*) | 0.00339 |
|  | Metabolomics | sphingomyelin (d18:1/25:0, d19:0/24:1, d20:1/23:0, d19:1/24:0)* | 0.0 |
|  |  | sulfate of piperine metabolite C16H19NO3 (2)* | 0.0 |
|  |  | suberoylcarnitine (C8-DC) | 0.0 |
|  |  | suberate (C8-DC) | 0.0 |
|  | Lipidomics | Unknown_mz_768.54962_- RT_23.304 | 0.0 |
|  |  | Unknown_mz_763.31421_+ RT_5.276 | 0.0 |
| Severe | Transcriptomics | C-C Motif Chemokine Ligand 4 (*CCL4*) | 0.00612 |
|  |  | C-X-C Motif Chemokine Ligand 1 (*CXCL1*) | 0.00549 |
|  | Proteomics | Interleukin 7 Receptor (*IL7R*) | 0.00358 |
|  | Metabolomics | sphingomyelin (d18:2/21:0, d16:2/23:0)* | 0.0 |
|  |  | tartronate (hydroxymalonate) | 0.0 |
|  |  | sulfate of piperine metabolite C18H21NO3 (1)* | 0.0 |
|  | Lipidomics | Unknown_mz_765.60468_+ RT_16.383 | 0.0 |
|  |  | Unknown_mz_766.75885_+ RT_1.234 | 0.0 |
|  |  | Unknown_mz_766.69232_+ RT_28.305 | 0.0 |

#### Random walk analysis on disease-state omics-specific graphs using hypothesis-driven seeds

Like the network exploration using data-driven seeds, we repeated the analysis with hypothesis-driven seeds. We observed from the multi-layer graphs (accessible at http://cytoscape.h3africa.org) that *IL-6* interaction with features of different omics data types increased with disease severity, thus indicating the differential association of *IL-6* with the different disease states. On the other hand, we observed *IL-6R* interactions mainly with proteins and transcripts to increase with disease severity as compared to interactions with metabolites. The results suggest *IL-6* interaction with proteins (e.g., *IFNB*, *IFIT3*), transcripts (e.g., *CXCL1*, *CXCL2, CCL3*), and metabolites (e.g., 1-(1-enyl-palmitoyl)-GPC, 1-(1-enyl-palmitoyl)-2-oleoyl-GPC (P-16:0/18:1)) may contribute to its significant role in disease severity.

### Evaluating features and interactions of generated multi-layered graphs

In this section, we dissect different multi-layered graphs generated from random walk analysis to examine common and unique feature interactions related to disease severity. The multi-layered networks generated for each disease state contain four types of nodes (proteins, transcripts, lipids, and metabolites), and at most six types of edges (transcript-transcript, protein-protein, protein-transcript, protein-metabolite, metabolite-metabolite, lipid-lipid). We did not observe protein-lipid edge types across all the networks. The seed exploration prioritizes nodes that have an either direct or indirect connection to the seeds, thus no observed protein-lipid edge type is an indication that there was limited bipartite data that captures interactions between nodes and seeds of interest.

### Evaluating multi-layered graphs generated using data-driven seeds

We evaluated the feature interactions (**Supplementary File 2**) present in the multi-layer graphs (accessible at http://cytoscape.h3africa.org) identifying 204 interactions associated with a specific disease state (**Figure 4A**), of which 79%, 15%, and 6% are transcript-transcript, lipid-lipid, and protein-protein interactions respectively. Of these interactions, most (88%) are associated with the mild disease state (**Figure 4A**). Specifically, we observed interactions between *CCL4* and human leucocyte antigen and other co-stimulatory molecules (e.g., *CD2, CD4, CD8, CD83, CD53*, *CD3D HLA-DPA1, HLA-DPB1, HLA-DRA*), and other molecules expressed in monocytes and macrophages (e.g., *CCL5, CCL7, CCL8*) to be highly predominant in the mild disease state. This observation may suggest a more efficient immune response to the virus among patients with mild disease as compared to those with moderate and severe disease, thus leading in the latter group to decreased recruitment of immune cells to the site of infection (30).

**Figure 4.**
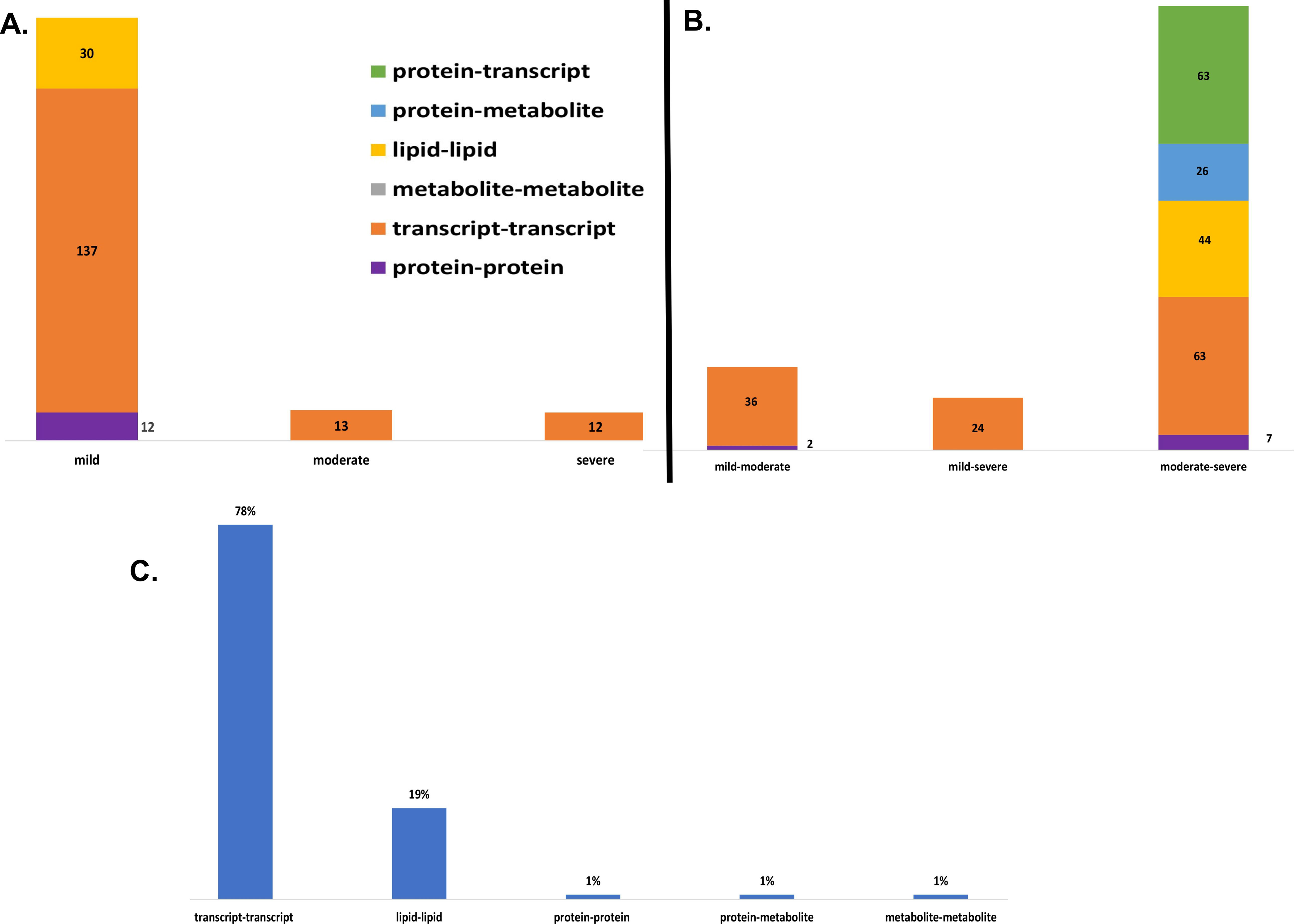
(**A**) Illustration of the distribution of interactions between omics features associated with one disease state identified from the data-driven approach (**B**) Illustration of the distribution of interactions between omics features associated with two disease states identified from the data-driven approach (**C**) Illustration of the distribution of interactions between omics features associated with three disease states identified from the data-driven approach

Additional investigation of 263 interactions associated with only two of the three disease states revealed more pairwise interactions common between the moderate and severe disease states as compared to those in common between the mild-moderate and mild-severe disease state pairs (**Figure 4B**).

We identified intra-omics interactions (specifically transcript-transcript and protein-protein) shared between mild-severe and mild-moderate disease state pairs. Notable interactions were among *STAT1,* inteferons (e.g., *IRF1, IRF6, IFNB1, ISG20*) and *SOD2*. *STAT1* is involved in regulating T-cell activation and differentiation responses, thus regulating the pathogenesis of COVID-19. Therefore, interactions between *STAT1* and T-cell receptors (e.g., *CD38, CD40, CD48, CD68*) in the mild-severe disease state, may suggest a more efficient role in the immune response to viral infections during mild disease state and higher *STAT1* activation in severe disease state contributing to the cytokine storm and hyperinflammation that are characteristic of severe COVID-19 (36). Further, interactions between *STA1* and interferons highlight the critical role of *STAT1* in the regulation of interferon-stimulated genes, because interferons are key cytokines in the immune response to viral infections. Also, interactions between interferons and other genes (e.g., *TNFRSF25*, *TLR3, MAPK1*) involved in interferon-mediated pathways, highlight the likelihood of innate and adaptive immune stimulatory effects.

A distinctive factor for severe-moderate compared to mild-moderate and mild-severe disease state pairs was the cross-layer interactions between protein-metabolites and protein-transcripts. Particularly, Hepatocyte Growth Factor (*HGF*) and Matrix Metallopeptidase 12 (*MMP12*) interactions with metabolites (e.g., 6-bromotryptophan, cortolone glucuronide, 1-palmityl-2-linoleoyl-GPC, 1-(1-enyl-stearoyl)-2-linoleoyl-GPC, sphingomyelin) and proteins implicated in various cellular processes were highly predominant.

Evidence suggests that both *HGF* and *MMP12* levels are significantly elevated in the lungs of patients with severe disease and play a role in disease pathogenesis and lung injury (37). Therefore, the observed cross-layer interactions for *HGF* and *MMP12* in moderate and severe diseases suggest that cross-layer interactions influence clinical heterogeneity, thus influencing the dynamics of disease severity. Furthermore, *HGF* has been shown to have antiviral effects against the SARS-CoV-2 virus *in vitro*, suggesting that it may help to inhibit viral replication in infected cells (38, 39). Thus, the observed interactions may suggest a protective role in ameliorating the progression of COVID-19, as well as a target for drug research (38, 39).

Also, an investigation of 397 interactions common across the three disease states revealed a higher proportion of interactions between transcripts associated with various cellular processes and about 1% each for metabolite-metabolite, protein-protein, and protein-metabolite interactions (**Figure 4C**).

#### Evaluating multi-layered graphs generated using hypothesis-driven seeds

We evaluated the feature interactions (**Supplementary File 2**) in the multi-layered graphs (accessible at http://cytoscape.h3africa.org) generated from hypothesis-driven seeds (**Table 1**). and identified 807 interactions associated with one disease state of which approximately 37%, 37%, 2%, 15%, and 9% are transcript-transcript, lipid-lipid, protein-protein, metabolite-metabolite, and protein-metabolite interactions respectively (**Figure 5A**). Also, of these interactions, approximately 20%, 40%, and 40% are associated with mild, moderate, and severe disease states, respectively. We identified for mild disease state, interactions between *CCL2* and interlukins (e.g., *IL-13, IL-18, IL-1R1, IL-1R1, IL-1R2, IL-6ST, IL-1B, IRF9*), chemokines (e.g., *CXCL10, CXCL11, CXCL9*, *CCL8*, *CCL7*), and other features involved in immune response. Notable cross-layer interactions for the mild disease state were between *CCL2* and metabolites involved in various metabolic and inflammatory processes, including but not limited to, taurine, 1-(1-enyl-palmitoyl)-2-oleoyl-GPC, 1-(1-enyl-palmitoyl)-GPC, 1-(1-enyl-palmitoyl)-GPE, 1-(1-enyl-stearoyl)-2-dihomo-linolenoyl-GPE, behenoyl sphingomyelin, 1-(1-enyl-palmitoyl)-GPC, 6-bromotryptophan, 1-margaroyl-GPC, and 1-myristoyl-GPC. This observation may further suggest a more efficient immune response to the virus among patients with mild disease.

**Figure 5.**
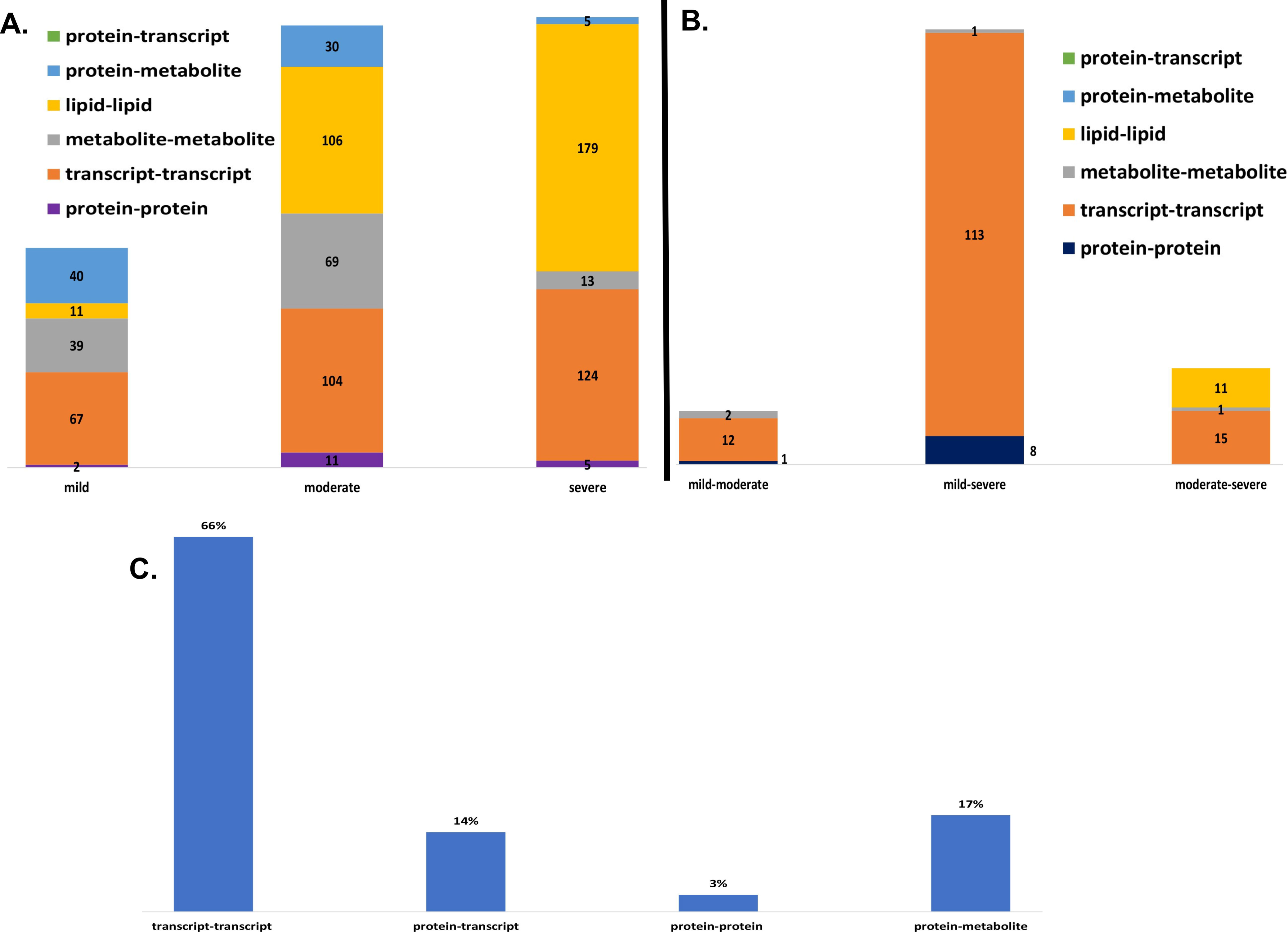
(**A**) Illustration of the distribution of interactions between omics features associated with one disease state identified from the hypothesis-driven approach (**B**) Illustration of the distribution of interactions between omics features associated with two disease states identified from the hypothesis-driven approach (**C**) Illustration of the distribution of interactions between omics features associated with three disease states identified from the hypothesis-driven approach

For the moderate and severe disease states, we identified subnetworks between *IL10, CXCL1,* and *NFKB1* with chemokines (*CXCL10, CXCL12, CXCL5, CXCL8*, *CCL3*) and other biosignatures involved in immune and metabolic processes. Notable cross-layer interactions were interactions between *IL10* and *IL-6* with metabolites. The metabolites including but not limited to 1-(1-enyl-palmitoyl)-2-oleoyl-GPC (P-16:0/18:1)*, 1-(1-enyl-palmitoyl)-GPC (P-16:0)*, 1-(1-enyl-palmitoyl)-GPE (P-16:0)*, 1-(1-enyl-stearoyl)-2-dihomo-linolenoyl-GPE (P-18:0/20:3)*, phosphatidylethanolamine, taurine, 1-(1-enyl-stearoyl)-GPE (P-18:0)*, and 11beta-hydroxyandrosterone glucuronide are involved in various metabolic and inflammatory processes. There is evidence of a correlation between *IL-6* levels and altered levels of various metabolites including amino acids, fatty acids, and lipids among severe COVID-19 patients (40, 41). Studies have also reported varying correlations between *IL-10* levels with altered metabolite levels in infected COVID-19 patients (41). For instance, one study found that in COVID-19 patients, *IL-10* levels were positively correlated with metabolites involved in glycolysis and the pentose phosphate pathway, such as glucose, fructose, and ribose-5-phosphate. The study also found a negative correlation between *IL-10* levels and metabolites involved in the tricarboxylic acid cycle (TCA cycle) and oxidative phosphorylation, such as citrate, succinate, and ATP. These findings suggest that *IL-10* may be associated with a shift in cellular metabolism towards glycolysis and away from oxidative phosphorylation in COVID-19 patients (41).

Furthermore, the analysis of interactions involved with only two disease states revealed 335 interactions, out of which approximately 6%, 85%, 7%, and 2% are protein-protein, transcript-transcript, lipid-lipid, and metabolite-metabolite interactions, respectively (**Figure 5B**). Of these interactions, approximately 16%, 10*%,* and 74% are involved with moderate-severe, mild-moderate, and mild-severe disease states, respectively.

Finally, we identified 894 interactions common across the three disease states, of which approximately 66%, 3%, 14%, and 17% are transcript-transcript, protein-protein, protein-transcript, and protein-metabolite interactions respectively (**Figure 5C**). We identified subnetworks mainly formed by *IL7R*, *IL-6*, and *IL-67.* Notable interactions formed by *IL7R*, *IL-6*, and *IL-67* were between *HLAs* (e.g., *HLA-A, HLA-B, HLA-C, HLA-DRA, HLA-DPA1, HLA-E, HLA-F*), interleukins (e.g., *IL-10, IL10RA, IL13, IL-16, IL-18, IL-1A, IL-1B, IL-1R2, IL-1RN, IL2, IL2RA, IL3RA*,) and other immune response-related genes (e.g., *ICAM1, FCER1G, FCGR3A, FCGR3B*), suggesting immune response to the virus.

We also identified cross-layer interactions between *IL-6*, and metabolites (e.g., glycerophosphoethanolamine, 1-adrenoyl-GPC, 1-margaroyl-GPC, 1-myristoyl-2-linoleoyl-GPC). The overall results highlight not only the influence of *IL-6* in COVID-19 but also suggested that cross-layer interactions involving IL6 influence clinical heterogeneity, thus influencing the dynamics of disease severity.

### Characterizing multi-layered graphs

The observation that cross-layer interactions appear to be a distinctive factor for moderate and severe disease states using both data- and hypothesis-driven methods necessitated the determination of network statistics to further characterize the multi-layered graphs.

According to network density, network heterogeneity, and characteristic path length statistical metrics, graphs with high characteristic route length values have high network heterogeneity and comparatively low network density (**Supplementary data**). The statistical analysis (**Table 4**) further supported the idea that cross-layer interactions could be a factor underlying heterogeneity in disease severity among patients.

**Table 4.**
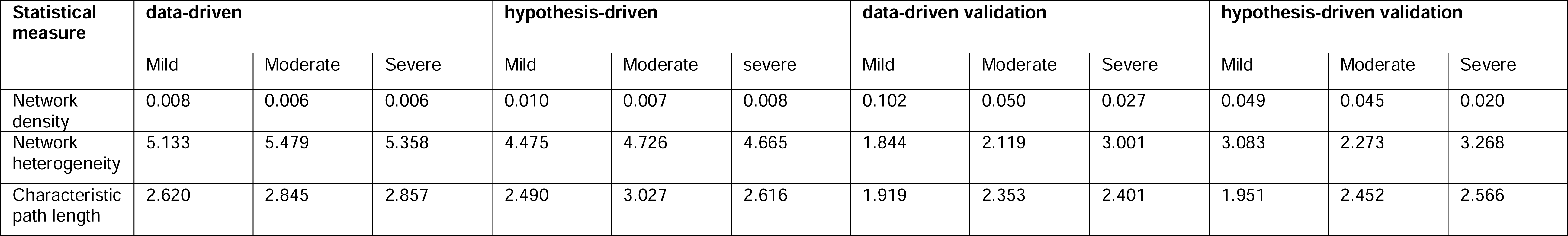
Results from the statistical analysis of generated multi-layered graphs.

### Identifying disease states biosignature

#### Biosignatures discriminating between disease states based on data-driven seeds

The results of the multi-layer analysis (**Supplementary File 2**) formed the basis for identifying features that discriminate between disease states. Specifically, we explored the pairwise relations associated with one disease state (**Supplementary File 2**), as determined using the data-driven approach, and identified 173 discriminatory features (**Table 5**). Of note, the features identified are likely involved in all disease and non-disease phases because they form part of the biological system. However, identifying these features to discriminate between disease states in our analysis may suggest that they are differentially associated (either up or down regulated) in specific disease states.

**Table 5.**
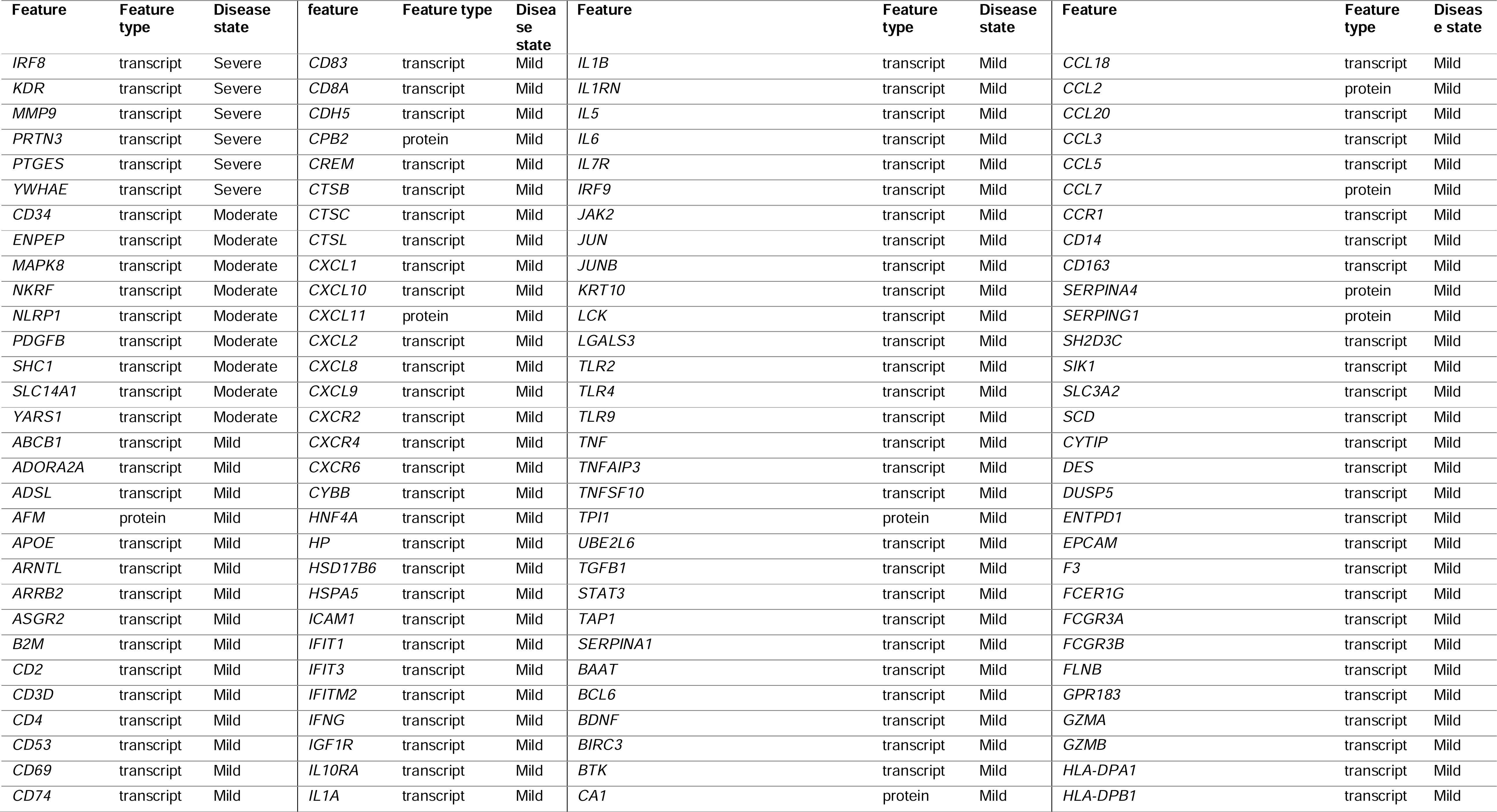
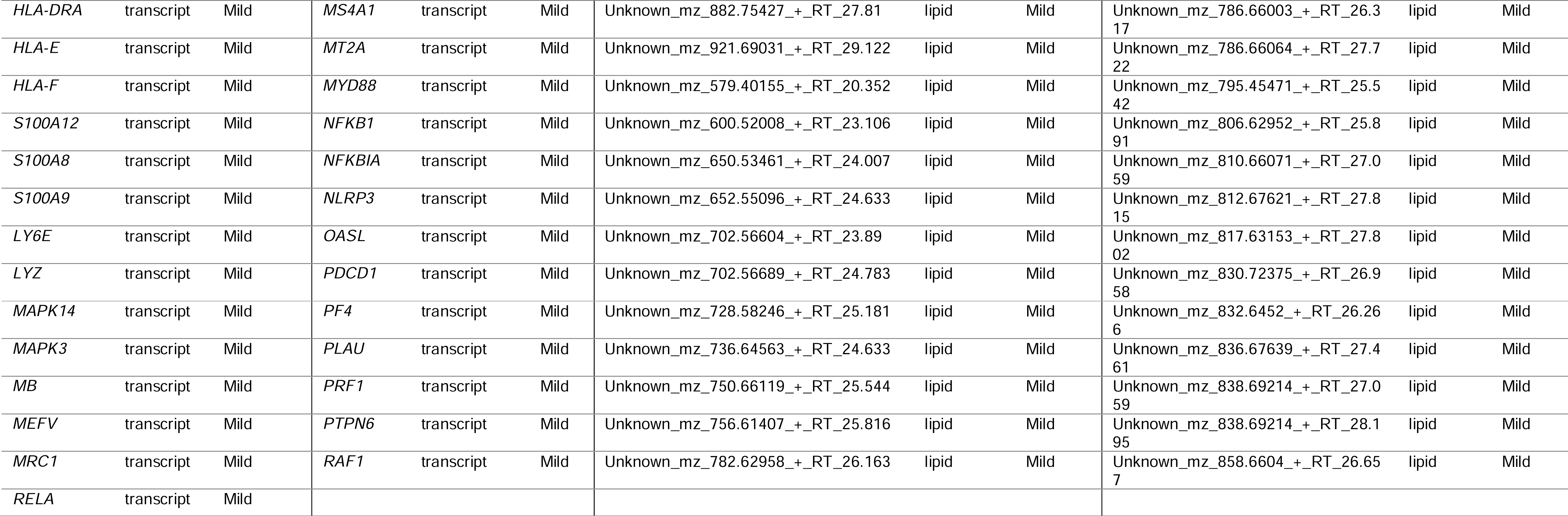
Identified biosignatures that discriminate disease states based on data-driven seeds.

Of the 158 discriminatory features that were differentially associated with the mild disease state, approximately 79%, 6%, and 16% were transcripts, proteins, and (uncharacterized) lipids respectively. We identified chemokines (e.g., *CXCL10, CXCL12, CXCL5, CXCL8*, *CCL3, CCL8*), T-cell receptors (e.g., *CD38, CD40, CD48, CD68*), *HLAs* (e.g., *HLA-DPA1, HLA-DPB1, HLA-DRA, HLA-E, HLA-F*), interferons (e.g, *IFIT1, IFIT3, IFITM2*), Toll-like receptors (e.g., *TLR2, TLR4, TLR9*) to discriminate the mild disease states. These features are involved in immune responses and play a part in viral entry into host T-cells (30, 42-44). For instance, *TLR2* activation increased the expression of ACE2, the receptor that SARS-CoV-2 uses to enter cells, suggesting that *TLR2* may play a role in viral entry into hosT-cells. *IFIT1* has been shown to have antiviral activity against SARS-CoV-2, and may thus be an important component of the body’s immune response against the virus (30). An elevated level of *CCL18* is associated with inflammation in the lungs of COVID-19 patients through the recruitment and activation of immune cells, including T-cells and dendritic cells in the lungs (45). Importantly, biosignatures including but not limited to HLA class I alleles, and *CXCL12*, have been validated through sequencing and cohort screening techniques to play a relevant role in immune defense against SARS-CoV-2 (46, 47).

Only 9 and 6 transcripts were associated with moderate and severe disease states respectively: no other omics features were identified to be associated with either of these disease states. There were also no metabolites that were differentially associated with any one of the disease states. This indicated that transcripts strongly differentiate the mild disease state from the moderate and severe disease states.

#### Biosignatures discriminating between disease states based on hypothesis-driven seeds

We explored the pairwise relations associated with one disease state (**Supplementary File 2)** based on analysis using seeds that were selected to test specific hypotheses. The results (**Table 6**) revealed more biosignatures to be differentially associated with moderate and severe disease states than the mild disease state. Additionally, unlike with the data-driven seed analysis, we observed more proteins and metabolites that discriminated between the moderate disease state than the others.

**Table 6.** Identified biosignatures that discriminate disease states based on hypothesis-driven seeds.

| Feature | Feature type | Disease state | Feature | Feature type | Disease state | Feature | Feature type | Disease state |
| --- | --- | --- | --- | --- | --- | --- | --- | --- |
| *ABCC1* | Transcript | Severe | Unknown_mz_742.65503+_RT_26.11 | Lipid | Severe | *TNF* | Transcript | Moderate |
| *ACKR3* | Transcript | Severe | Unknown_mz_743.61646+_RT_28.297 | Lipid | Severe | *TNFRSF1A* | Transcript | Moderate |
| *ACOD1* | Transcript | Severe | Unknown_mz_745.5152+_RT_14.3 | Lipid | Severe | *TXN* | Transcript | Moderate |
| *ADRA2B* | Transcript | Severe | Unknown_mz_748.39844-_RT_24.778 | Lipid | Severe | *TYRO3* | Transcript | Moderate |
| *ATP6AP2* | Transcript | Severe | Unknown_mz_752.67621+_RT_29.183 | Lipid | Severe | *UBE3A* | Transcript | Moderate |
| *BCL6* | Transcript | Severe | Unknown_mz_752.67676+_RT_26.296 | Lipid | Severe | *UNG* | Transcript | Moderate |
| *BIRC3* | Transcript | Severe | Unknown_mz_757.63165+_RT_28.819 | Lipid | Severe | *VWF* | Transcript | Moderate |
| *CA1* | Transcript | Severe | Unknown_mz_762.66071+_RT_27.74 | Lipid | Severe | *AREG* | Protein | Moderate |
| *CA2* | Transcript | Severe | Unknown_mz_764.67676+_RT_27.522 | Lipid | Severe | *EPO* | Protein | Moderate |
| *CCL16* | Transcript | Severe | Unknown_mz_765.60468+_RT_16.383 | Lipid | Severe | *IL17A* | Protein | Moderate |
| *CD69* | Transcript | Severe | Unknown_mz_766.69196+_RT_26.293 | Lipid | Severe | *NGF* | Protein | Moderate |
| *CD83* | Transcript | Severe | Unknown_mz_766.69232+_RT_28.305 | Lipid | Severe | *THPO* | Protein | Moderate |
| *CFP* | Transcript | Severe | Unknown_mz_769.6311+_RT_28.523 | Lipid | Severe | *TNNI3* | Protein | Moderate |
| *CREM* | Transcript | Severe | Unknown_mz_770.68744+_RT_27.138 | Lipid | Severe | *VEGFA* | Protein | Moderate |
| *CSF3* | Transcript | Severe | Unknown_mz_771.64661+_RT_29.293 | Lipid | Severe | Unknown_mz_404.19128+_RT_4.73 | Lipid | Moderate |
| *CXCL2* | Transcript | Severe | Unknown_mz_776.67676+_RT_28.157 | Lipid | Severe | Unknown_mz_405.13147-_RT_5.625 | Lipid | Moderate |
| *CXCL3* | Transcript | Severe | Unknown_mz_780.69897+_RT_28.309 | Lipid | Severe | Unknown_mz_405.13184-_RT_1.471 | Lipid | Moderate |
| *CXCR1* | Transcript | Severe | Unknown_mz_780.7074+_RT_29.787 | Lipid | Severe | Unknown_mz_407.12833-_RT_5.279 | Lipid | Moderate |
| *CXCR2* | Transcript | Severe | Unknown_mz_780.7077+_RT_29.279 | Lipid | Severe | Unknown_mz_409.14612+_RT_4.35 | Lipid | Moderate |
| *CXCR3* | Transcript | Severe | Unknown_mz_780.70953+_RT_27.345 | Lipid | Severe | Unknown_mz_412.14255+_RT_6.178 | Lipid | Moderate |
| *CXCR6* | Transcript | Severe | Unknown_mz_782.4267-_RT_24.225 | Lipid | Severe | Unknown_mz_413.12015+_RT_6.629 | Lipid | Moderate |
| *DPP4* | Transcript | Severe | Unknown_mz_782.7207+_RT_29.713 | Lipid | Severe | Unknown_mz_415.12424-_RT_5.789 | Lipid | Moderate |
| *ELF3* | Transcript | Severe | Unknown_mz_792.70734+_RT_28.536 | Lipid | Severe | Unknown_mz_415.12424-_RT_6.059 | Lipid | Moderate |
| *EPX* | Transcript | Severe | Unknown_mz_793.63165+_RT_28.367 | Lipid | Severe | Unknown_mz_415.14279-_RT_6.442 | Lipid | Moderate |
| *GCLC* | Transcript | Severe | Unknown_mz_794.72351+_RT_29.303 | Lipid | Severe | Unknown_mz_418.17029+_RT_4.744 | Lipid | Moderate |
| *GJA1* | Transcript | Severe | Unknown_mz_794.7243+_RT_28.327 | Lipid | Severe | Unknown_mz_418.17029+_RT_5.022 | Lipid | Moderate |
| *IDO1* | Transcript | Severe | Unknown_mz_795.64679+_RT_28.628 | Lipid | Severe | Unknown_mz_419.20432-_RT_6.987 | Lipid | Moderate |
| *IFITM1* | Transcript | Severe | Unknown_mz_796.73724+_RT_30.218 | Lipid | Severe | Unknown_mz_431.11887-_RT_4.204 | Lipid | Moderate |
| *IFITM2* | Transcript | Severe | Unknown_mz_798.7207+_RT_28.176 | Lipid | Severe | Unknown_mz_432.15057-_RT_5.265 | Lipid | Moderate |
| *IL15* | Transcript | Severe | Unknown_mz_806.72339+_RT_29.498 | Lipid | Severe | Unknown_mz_433.11655-_RT_7.027 | Lipid | Moderate |
| *IL33* | Transcript | Severe | Unknown_mz_811.62109+_RT_28.63 | Lipid | Severe | Unknown_mz_437.14285-_RT_5.276 | Lipid | Moderate |
| *IL3RA* | Transcript | Severe | Unknown_mz_812.67621+_RT_28.125 | Lipid | Severe | Unknown_mz_440.13779-_RT_5.443 | Lipid | Moderate |
| *IL7R* | Transcript | Severe | Unknown_mz_814.69196+_RT_28.152 | Lipid | Severe | Unknown_mz_449.11169-_RT_4.82 | Lipid | Moderate |
| *IRF8* | Transcript | Severe | Unknown_mz_816.7077+_RT_29.317 | Lipid | Severe | Unknown_mz_453.08536-_RT_7.227 | Lipid | Moderate |
| *ITGAV* | Transcript | Severe | Unknown_mz_820.73932+_RT_28.552 | Lipid | Severe | Unknown_mz_458.14764+_RT_5.412 | Lipid | Moderate |
| *ITGAX* | Transcript | Severe | Unknown_mz_821.66254+_RT_28.831 | Lipid | Severe | Unknown_mz_471.09415+_RT_5.274 | Lipid | Moderate |
| *ITGB3* | Transcript | Severe | Unknown_mz_822.72083+_RT_27.527 | Lipid | Severe | Unknown_mz_476.08188-_RT_6.986 | Lipid | Moderate |
| *JUN* | Transcript | Severe | Unknown_mz_822.75537+_RT_28.304 | Lipid | Severe | Unknown_mz_494.01953-_RT_6.498 | Lipid | Moderate |
| *KDM5B* | Transcript | Severe | Unknown_mz_824.737+_RT_28.3 | Lipid | Severe | Unknown_mz_494.02023-_RT_5.961 | Lipid | Moderate |
| *KDR* | Transcript | Severe | Unknown_mz_826.75671+_RT_29.2 | Lipid | Severe | Unknown_mz_496.01746-_RT_5.962 | Lipid | Moderate |
| *KIT* | Transcript | Severe | Unknown_mz_830.71057+_RT_27.868 | Lipid | Severe | Unknown_mz_506.16876+_RT_4.424 | Lipid | Moderate |
| *KITLG* | Transcript | Severe | Unknown_mz_832.72412+_RT_28.634 | Lipid | Severe | Unknown_mz_511.16394-_RT_5.692 | Lipid | Moderate |
| *KRT5* | Transcript | Severe | Unknown_mz_832.73975+_RT_27.702 | Lipid | Severe | Unknown_mz_512.05823-_RT_6.993 | Lipid | Moderate |
| *LPAR2* | Transcript | Severe | Unknown_mz_836.74402+_RT_28.521 | Lipid | Severe | Unknown_mz_514.05322-_RT_6.984 | Lipid | Moderate |
| *LRP1* | Transcript | Severe | Unknown_mz_836.77008+_RT_29.296 | Lipid | Severe | Unknown_mz_529.19226-_RT_7.004 | Lipid | Moderate |
| *MAF* | Transcript | Severe | Unknown_mz_846.75452+_RT_27.716 | Lipid | Severe | Unknown_mz_529.99689-_RT_5.25 | Lipid | Moderate |
| *MAPK14* | Transcript | Severe | Unknown_mz_846.7561+_RT_28.63 | Lipid | Severe | Unknown_mz_548.2337+_RT_7.003 | Lipid | Moderate |
| *MCAM* | Transcript | Severe | Unknown_mz_850.7865+_RT_29.293 | Lipid | Severe | Unknown_mz_551.17395-_RT_6.21 | Lipid | Moderate |
| *MDM2* | Transcript | Severe | Unknown_mz_851.5166+_RT_27.535 | Lipid | Severe | Unknown_mz_553.18915+_RT_6.211 | Lipid | Moderate |
| *ME1* | Transcript | Severe | Unknown_mz_852.76886+_RT_29.292 | Lipid | Severe | Unknown_mz_557.16913-_RT_5.138 | Lipid | Moderate |
| *MMP1* | Transcript | Severe | Unknown_mz_862.78735+_RT_29.482 | Lipid | Severe | Unknown_mz_573.16394-_RT_5.023 | Lipid | Moderate |
| *MMP2* | Transcript | Severe | Unknown_mz_864.77179+_RT_28.55 | Lipid | Severe | Unknown_mz_643.24255-_RT_5.275 | Lipid | Moderate |
| *MSN* | Transcript | Severe | Unknown_mz_876.77521+_RT_28.63 | Lipid | Severe | Unknown_mz_643.24255-_RT_5.626 | Lipid | Moderate |
| *MSR1* | Transcript | Severe | Unknown_mz_902.78906+_RT_28.837 | Lipid | Severe | Unknown_mz_659.23743-_RT_5.596 | Lipid | Moderate |
| *MT2A* | Transcript | Severe | Unknown_mz_905.5639+_RT_28.63 | Lipid | Severe | Unknown_mz_659.23773-_RT_5.268 | Lipid | Moderate |
| *OPRD1* | Transcript | severe | Unknown_mz_1204.04907+_RT_24.162 | Lipid | severe | Unknown_mz_707.32788-_RT_7.231 | Lipid | Moderate |
| *OPRM1* | Transcript | severe | Unknown_mz_1227.05237+_RT_24.126 | Lipid | severe | Unknown_mz_739.31799-_RT_5.276 | Lipid | Moderate |
| *OSM* | Transcript | severe | Unknown_mz_1237.12878+_RT_25.831 | Lipid | severe | Unknown_mz_753.29736-_RT_5.638 | Lipid | Moderate |
| *PLAU* | Transcript | severe | Unknown_mz_1455.19128+_RT_24.513 | Lipid | severe | Unknown_mz_760.24139-_RT_7.23 | Lipid | Moderate |
| *PLXNA2* | Transcript | severe | Unknown_mz_1468.27539+_RT_29.197 | Lipid | severe | Unknown_mz_761.30078-_RT_5.276 | Lipid | Moderate |
| *PTGS1* | Transcript | severe | Unknown_mz_1494.28638+_RT_29.248 | Lipid | severe | Unknown_mz_763.31421+_RT_5.276 | Lipid | Moderate |
| *RELA* | Transcript | severe | Unknown_mz_1516.27661+_RT_28.532 | Lipid | severe | Unknown_mz_768.54962-_RT_23.304 | Lipid | Moderate |
| *RNASE1* | Transcript | severe | Unknown_mz_1567.38342+_RT_29.458 | Lipid | severe | Unknown_mz_785.24414+_RT_7.219 | Lipid | Moderate |
| *RTN2* | Transcript | severe | Unknown_mz_1572.33655+_RT_29.463 | Lipid | severe | Unknown_mz_822.26886-_RT_5.611 | Lipid | Moderate |
| *S100A4* | Transcript | severe | Unknown_mz_245.22595+_RT_13.471 | Lipid | severe | Unknown_mz_854.29572-_RT_5.701 | Lipid | Moderate |
| *S100A9* | Transcript | severe | Unknown_mz_258.11005+_RT_1.474 | Lipid | severe | Unknown_mz_864.41992-_RT_7.232 | Lipid | Moderate |
| *S1PR1* | Transcript | severe | Unknown_mz_263.23645+_RT_13.471 | Lipid | severe | Unknown_mz_886.25458-_RT_5.207 | Lipid | Moderate |
| *SDC2* | Transcript | severe | Unknown_mz_287.23691+_RT_13.446 | Lipid | severe | Unknown_mz_904.3288-_RT_5.061 | Lipid | Moderate |
| *SDCBP* | Transcript | severe | Unknown_mz_302.10052-_RT_1.27 | Lipid | severe | PC 33:3_RT_22.262 | Lipid | Moderate |
| *SELENOP* | Transcript | severe | Unknown_mz_311.21869+_RT_10.981 | Lipid | severe | PE 16:0_18:1_RT_22.866 | Lipid | Moderate |
| *SLC14A1* | Transcript | severe | Unknown_mz_312.29016+_RT_17.443 | Lipid | severe | PE 18:1_18:1_RT_20.883 | Lipid | Moderate |
| *SLC1A5* | Transcript | severe | Unknown_mz_313.27353+_RT_14.323 | Lipid | severe | PE 18:2_18:1_RT_22.266 | Lipid | Moderate |
| *SLC2A1* | Transcript | severe | Unknown_mz_317.24741+_RT_12.26 | Lipid | severe | PE 35:1_RT_23.425 | Lipid | Moderate |
| *SOD2* | Transcript | severe | Unknown_mz_319.22717-_RT_11.732 | Lipid | severe | PE 36:2_RT_23.241 | Lipid | Moderate |
| *SP1* | Transcript | severe | Unknown_mz_337.27298+_RT_13.471 | Lipid | severe | PE 38:5_RT_22.233 | Lipid | Moderate |
| *STT3A* | Transcript | severe | Unknown_mz_339.25079+_RT_13.262 | Lipid | severe | Plasmenyl-PE P-18:0_18:2_RT_22.986 | Lipid | Moderate |
| *TCN1* | Transcript | severe | Unknown_mz_726.63995+_RT_27.125 | Lipid | severe | Unknown_mz_1066.38086-_RT_5.093 | Lipid | Moderate |
| *TNFSF10* | Transcript | severe | Unknown_mz_733.57343+_RT_28.179 | Lipid | severe | Unknown_mz_205.08609+_RT_4.89 | Lipid | Moderate |
| *VIPR1* | Transcript | severe | Unknown_mz_734.63037+_RT_25.901 | Lipid | severe | Unknown_mz_213.96338-_RT_4.886 | Lipid | Moderate |
| *ZFP36* | Transcript | severe | Unknown_mz_736.64594+_RT_26.57 | Lipid | severe | Unknown_mz_223.09703-_RT_6.088 | Lipid | Moderate |
| *PGF* | Protein | severe | Unknown_mz_738.53009+_RT_13.467 | Lipid | severe | Unknown_mz_239.09181-_RT_4.887 | Lipid | Moderate |
| *STC1* | Protein | severe | Unknown_mz_738.6001+_RT_14.757 | Lipid | severe | Unknown_mz_239.09184-_RT_5.113 | Lipid | Moderate |
| 1-palmitoyl-2-arachidonoyl-GPE (16:0/20:4)* | Metabolite | severe | (S)-a-amino-omega-caprolactam | Metabolite | moderate | Unknown_mz_269.06628-_RT_4.558 | Lipid | Moderate |
| choline phosphate | Metabolite | Severe | 11beta-hydroxyandrosterone glucuronide | Metabolite | moderate | Unknown_mz_273.07901+_RT_5.853 | Lipid | Moderate |
| glycerate | Metabolite | Severe | 11-ketoetiocholanolone glucuronide | Metabolite | moderate | Unknown_mz_287.05914-_RT_5.49 | Lipid | Moderate |
| oxalate (ethanedioate) | Metabolite | Severe | 1-arachidonoyl-GPE (20:4n6)* | Metabolite | moderate | Unknown_mz_289.07495-_RT_4.915 | Lipid | Moderate |
| sphingomyelin (d17:2/16:0, d18:2/15:0)* | Metabolite | Severe | 1-linoleoyl-GPE (18:2)* | Metabolite | moderate | Unknown_mz_303.05392-_RT_4.963 | Lipid | Moderate |
| sphingomyelin (d18:2/14:0, d18:1/14:1)* | Metabolite | Severe | 1-methyl-4-imidazoleacetate | Metabolite | moderate | Unknown_mz_308.11621+_RT_4.916 | Lipid | Moderate |
| sphingomyelin (d18:2/18:1)* | Metabolite | Severe | 1-ribosyl-imidazoleacetate* | Metabolite | moderate | Unknown_mz_308.11621+_RT_5.265 | Lipid | Moderate |
| sphingomyelin (d18:2/21:0, d16:2/23:0)* | Metabolite | Severe | 2,3-dihydroxy-5-methylthio-4-pentenoate (DMTPA)* | Metabolite | moderate | Unknown_mz_315.19626-_RT_6.994 | Lipid | Moderate |
| sphingomyelin (d18:2/23:1)* | Metabolite | Severe | 3-(3-amino-3-carboxypropyl)uridine* | Metabolite | moderate | Unknown_mz_317.13824+_RT_5.979 | Lipid | Moderate |
| sulfate of piperine metabolite C18H21NO3 (1)* | Metabolite | Severe | 5-(galactosylhydroxy)-L-lysine | Metabolite | moderate | Unknown_mz_319.0676-_RT_6.574 | Lipid | Moderate |
| tartronate (hydroxymalonate) | Metabolite | Severe | adipoylcarnitine (C6-DC) | Metabolite | moderate | Unknown_mz_333.20648-_RT_6.997 | Lipid | Moderate |
| threonate | Metabolite | Severe | arabonate/xylonate | Metabolite | moderate | Unknown_mz_349.95889-_RT_6.159 | Lipid | Moderate |
| DG 12:0_14:0_RT_21.212 | Lipid | Severe | cortolone glucuronide (1) | Metabolite | moderate | Unknown_mz_351.13715+_RT_7.772 | Lipid | Moderate |
| LysoPC 18:2_RT_11.045 | Lipid | Severe | cystathionine | Metabolite | moderate | Unknown_mz_352.12076+_RT_6.347 | Lipid | Moderate |
| LysoPC 18:2_RT_11.623 | Lipid | Severe | decadienedioic acid (C10:2-DC)** | Metabolite | moderate | Unknown_mz_360.0907+_RT_5.792 | Lipid | Moderate |
| LysoPC 20:2_RT_12.854 | Lipid | Severe | erythronate* | Metabolite | moderate | Unknown_mz_361.94125-_RT_5.813 | Lipid | Moderate |
| LysoPC 20:3_RT_11.724 | Lipid | Severe | gamma-CEHC glucuronide* | Metabolite | moderate | Unknown_mz_361.9415-_RT_5.724 | Lipid | Moderate |
| LysoPC 20:3_RT_11.872 | Lipid | Severe | GlcNAc sulfate conjugate of C21H34O2 steroid** | Metabolite | moderate | Unknown_mz_368.1171-_RT_6.349 | Lipid | Moderate |
| LysoPC 20:3_RT_12.285 | Lipid | Severe | glucuronate | Metabolite | moderate | Unknown_mz_369.15408+_RT_5.026 | Lipid | Moderate |
| LysoPC 20:4_RT_10.912 | Lipid | Severe | glucuronide of piperine metabolite C17H21NO3 (3)* | Metabolite | moderate | Unknown_mz_369.15408+_RT_5.544 | Lipid | Moderate |
| LysoPC 20:4_RT_11.056 | Lipid | Severe | glucuronide of piperine metabolite C17H21NO3 (4)* | Metabolite | moderate | Unknown_mz_372.20126+_RT_7.226 | Lipid | Moderate |
| LysoPC 22:4_RT_12.289 | Lipid | Severe | glucuronide of piperine metabolite C17H21NO3 (5)* | Metabolite | moderate | Unknown_mz_375.21359+_RT_6.99 | Lipid | Moderate |
| LysoPC 22:4_RT_12.605 | Lipid | Severe | glutamine conjugate of C6H10O2 (2)* | Metabolite | moderate | Unknown_mz_375.21695-_RT_7.075 | Lipid | Moderate |
| LysoPC 22:5_RT_11.654 | Lipid | Severe | gulonate* | Metabolite | moderate | Unknown_mz_383.13434-_RT_6.069 | Lipid | Moderate |
| LysoPC 22:5_RT_12.087 | Lipid | Severe | hydantoin-5-propionate | Metabolite | moderate | Unknown_mz_385.14993-_RT_5.135 | Lipid | Moderate |
| LysoPC 22:6_RT_10.886 | Lipid | Severe | hydroxy-N6,N6,N6-trimethyllysine* | Metabolite | moderate | Unknown_mz_385.14999-_RT_1.473 | Lipid | Moderate |
| LysoPI 16:0_RT_11.045 | Lipid | Severe | hydroxypalmitoyl sphingomyelin (d18:1/16:0(OH))** | Metabolite | moderate | Unknown_mz_386.18082+_RT_2.416 | Lipid | Moderate |
| TG 10:0_15:0_16:0_RT_28.707 | Lipid | Severe | kynurenate | Metabolite | moderate | Unknown_mz_387.16437+_RT_4.35 | Lipid | Moderate |
| TG 12:0_12:0_14:0_RT_27.15 | Lipid | Severe | myo-inositol | Metabolite | moderate | Unknown_mz_387.19397_-_RT_6.987 | Lipid | Moderate |
| TG 12:0_12:0_18:2_RT_27.539 | Lipid | Severe | N,N-dimethyl-pro-pro | Metabolite | moderate | Unknown_mz_388.19608+_RT_5.18 | Lipid | Moderate |
| TG 12:0_12:0_18:3_RT_26.893 | Lipid | Severe | N2-acetyl,N6,N6-dimethyllysine | Metabolite | moderate | Unknown_mz_393.15167+_RT_6.505 | Lipid | Moderate |
| TG 12:0_16:1_16:1_RT_28.529 | Lipid | Severe | N6-carbamoylthreonyladenosine | Metabolite | moderate | Unknown_mz_397.91782_-_RT_5.723 | Lipid | Moderate |
| TG 16:0_16:0_24:0_RT_35.146 | Lipid | Severe | N6-succinyladenosine | Metabolite | moderate | Unknown_mz_397.91791_-_RT_5.811 | Lipid | Moderate |
| TG 16:0_18:0_18:0_RT_34.031 | Lipid | Severe | N-acetylkynurenine (2) | Metabolite | moderate | Unknown_mz_399.12878_-_RT_5.738 | Lipid | Moderate |
| TG 42:1_RT_28.302 | Lipid | Severe | N-acetyltryptophan | Metabolite | moderate | Unknown_mz_399.12924_-_RT_5.023 | Lipid | Moderate |
| TG 44:1_RT_29.294 | Lipid | Severe | O-sulfo-L-tyrosine | Metabolite | moderate | Unknown_mz_399.13174_-_RT_4.738 | Lipid | Moderate |
| TG 46:1_RT_30.294 | Lipid | Severe | p-cresol glucuronide* | Metabolite | moderate | Unknown_mz_399.91483_-_RT_5.811 | Lipid | Moderate |
| TG 46:2_RT_29.48 | Lipid | Severe | phenylacetylglutamine | Metabolite | moderate | Unknown_mz_401.14484_-_RT_3.792 | Lipid | Moderate |
| TG 46:3_RT_28.628 | Lipid | Severe | phenylalanylphenylalanine | Metabolite | moderate | Unknown_mz_402.17587+_RT_6.068 | Lipid | Moderate |
| TG 47:2_RT_29.947 | Lipid | Severe | pimeloylcarnitine/3-methyladipoylcarnitine (C7-DC) | Metabolite | moderate | Unknown_mz_404.19128+_RT_4.353 | Lipid | Moderate |
| TG 8:0_15:0_16:0_RT_27.751 | Lipid | Severe | piperine | Metabolite | moderate | diacylglycerol (16:1/18:2 [2], 16:0/18:3 [1])* | Metabolite | Mild |
| Unknown_mz_1039.67261+_RT_11.185 | Lipid | Severe | pseudouridine | Metabolite | moderate | dihomo-linolenoyl-choline | Metabolite | Mild |
| Unknown_mz_1041.68823+_RT_11.863 | Lipid | Severe | pyroglutamylvaline | Metabolite | moderate | docosahexaenoylcholine | Metabolite | Mild |
| Unknown_mz_1041.68872+_RT_11.738 | Lipid | Severe | quinolinate | Metabolite | moderate | dodecenedioate (C12:1-DC)* | Metabolite | Mild |
| Unknown_mz_1063.672+_RT_11.885 | Lipid | Severe | retinol (Vitamin A) | Metabolite | moderate | hexadecenedioate (C16:1-DC)* | Metabolite | Mild |
| Unknown_mz_1063.67212+_RT_11.077 | Lipid | Severe | S-adenosylhomocysteine (SAH) | Metabolite | moderate | lignoceroyl sphingomyelin (d18:1/24:0) | Metabolite | Mild |
| Unknown_mz_1063.67285+_RT_11.193 | Lipid | Severe | sphingomyelin (d18:1/17:0, d17:1/18:0, d19:1/16:0) | Metabolite | moderate | linoleoylcholine* | Metabolite | Mild |
| Unknown_mz_1087.69226_-_RT_12.293 | Lipid | Severe | sphingomyelin (d18:1/25:0, d19:0/24:1, d20:1/23:0, d19:1/24:0)* | Metabolite | moderate | oleoylcholine | Metabolite | Mild |
| Unknown_mz_1107.66125_-_RT_11.066 | Lipid | Severe | suberoylcarnitine (C8-DC) | Metabolite | moderate | palmitoylcholine | Metabolite | Mild |
| Unknown_mz_1109.65356+_RT_11.17 | Lipid | Severe | sulfate of piperine metabolite C16H19NO3 (2)* | Metabolite | moderate | phosphoethanolamine | Metabolite | Mild |
| Unknown_mz_1131.66211_-_RT_11.186 | Lipid | Severe | tetrahydrocortisol glucuronide | Metabolite | moderate | sphingomyelin (d18:1/14:0, d16:1/16:0)* | Metabolite | Mild |
| Unknown_mz_1155.04663+_RT_24.774 | Lipid | Severe | trans-4-hydroxyproline | Metabolite | moderate | stearoylcholine* | Metabolite | Mild |
| Unknown_mz_1181.06323+_RT_24.803 | Lipid | Severe | ADAM17 | Transcript | moderate | suberate (C8-DC) | Metabolite | Mild |
| Unknown_mz_1181.98779+_RT_23.981 | Lipid | Severe | AR | Transcript | moderate | tetradecanedioate (C14-DC) | Metabolite | Mild |
| Unknown_mz_1193.98499+_RT_23.882 | Lipid | Severe | ARRB2 | Transcript | moderate | tricosanoyl sphingomyelin (d18:1/23:0)* | Metabolite | Mild |
| Unknown_mz_339.28937_+_RT_14.753 | Lipid | Severe | *B2M* | Transcript | moderate | Unknown_mz_589.30219_+_RT_9.071 | Lipid | Mild |
| Unknown_mz_348.07925_+_RT_1.27 | Lipid | Severe | *BCL2A1* | Transcript | moderate | Unknown_mz_589.30377_-_RT_8.717 | Lipid | Mild |
| Unknown_mz_353.26541_+_RT_14.085 | Lipid | Severe | *CASP1* | Transcript | moderate | Unknown_mz_603.31915_-_RT_7.941 | Lipid | Mild |
| Unknown_mz_355.28354_+_RT_13.472 | Lipid | Severe | *CD19* | Transcript | moderate | Unknown_mz_721.54974_-_RT_20.015 | Lipid | Mild |
| Unknown_mz_357.29984_+_RT_14.753 | Lipid | Severe | *CD46* | Transcript | moderate | Unknown_mz_765.3349_-_RT_6.822 | Lipid | Mild |
| Unknown_mz_360.28915_+_RT_17.001 | Lipid | Severe | *CD8B* | Transcript | moderate | Unknown_mz_765.3349_+_RT_6.842 | Lipid | Mild |
| Unknown_mz_362.30511_+_RT_17.721 | Lipid | Severe | *CSF2* | Transcript | moderate | Unknown_mz_765.5752_-_RT_20.235 | Lipid | Mild |
| Unknown_mz_362.3056_+_RT_18.15 | Lipid | Severe | *EPCAM* | Transcript | moderate | Unknown_mz_767.35071_-_RT_6.389 | Lipid | Mild |
| Unknown_mz_365.24597_-_RT_14.058 | Lipid | Severe | *F10* | Transcript | moderate | Unknown_mz_769.36639_+_RT_6.393 | Lipid | Mild |
| Unknown_mz_369.23981_+_RT_14.301 | Lipid | Severe | *F8* | Transcript | moderate | Unknown_mz_794.50909_-_RT_19.3 | Lipid | Mild |
| Unknown_mz_370.29538_+_RT_12.394 | Lipid | Severe | *FCGR3A* | Transcript | moderate | Unknown_mz_935.31989_+_RT_6.496 | Lipid | Mild |
| Unknown_mz_372.31085_+_RT_13.272 | Lipid | Severe | *HIF1A* | Transcript | moderate | *CTLA4* | Transcript | Mild |
| Unknown_mz_372.31091_+_RT_13.953 | Lipid | Severe | *HMGB1* | Transcript | moderate | *DDIT3* | Transcript | Mild |
| Unknown_mz_375.25015_+_RT_12.546 | Lipid | Severe | *HSP90AA1* | Transcript | moderate | *ERG* | Transcript | Mild |
| Unknown_mz_377.26541_+_RT_12.394 | Lipid | Severe | *HSPA5* | Transcript | moderate | *FLT1* | Transcript | Mild |
| Unknown_mz_379.28143_+_RT_14.557 | Lipid | Severe | *IFIT3* | Transcript | moderate | *HAVCR2* | Transcript | Mild |
| Unknown_mz_379.28595_+_RT_13.314 | Lipid | Severe | *IFNAR2* | Transcript | moderate | *HLA-DRA* | Transcript | Mild |
| Unknown_mz_386.3259_+_RT_14.491 | Lipid | Severe | *IFNG* | Transcript | moderate | *IL1R2* | Transcript | Mild |
| Unknown_mz_391.24271_-_RT_13.475 | Lipid | Severe | *IFNGR2* | Transcript | moderate | *IL6ST* | Transcript | Mild |
| Unknown_mz_393.29718_+_RT_15.809 | Lipid | Severe | *IGF2R* | Transcript | moderate | *IRF9* | Transcript | Mild |
| Unknown_mz_396.31009_+_RT_13.295 | Lipid | Severe | *IKBKG* | Transcript | moderate | *ITGB2* | Transcript | Mild |
| Unknown_mz_396.31088_+_RT_13.446 | Lipid | Severe | *IL10RB* | Transcript | moderate | *LAMP3* | Transcript | Mild |
| Unknown_mz_400.34274_+_RT_15.344 | Lipid | Severe | *IL16* | Transcript | moderate | *MAOA* | Transcript | Mild |
| Unknown_mz_405.22116_+_RT_11.052 | Lipid | Severe | *IL1A* | Transcript | moderate | *MAS1* | Transcript | Mild |
| Unknown_mz_413.24588_-_RT_13.454 | Lipid | Severe | *IL2* | Transcript | moderate | *MUC1* | Transcript | Mild |
| Unknown_mz_420.31012_+_RT_13.354 | Lipid | Severe | *IL6R* | Transcript | moderate | *MX1* | Transcript | Mild |
| Unknown_mz_422.32593_+_RT_13.926 | Lipid | Severe | *IRF3* | Transcript | moderate | *NCAM1* | Transcript | Mild |
| Unknown_mz_445.2529_+_RT_13.463 | Lipid | Severe | *KCNJ2* | Transcript | moderate | *NLRP3* | Transcript | Mild |
| Unknown_mz_449.28555_+_RT_16.376 | Lipid | Severe | *LCK* | Transcript | moderate | *P2RX7* | Transcript | Mild |
| Unknown_mz_453.29675_+_RT_14.74 | Lipid | Severe | *MAPK1* | Transcript | moderate | *PLAAT4* | Transcript | Mild |
| Unknown_mz_496.37521_+_RT_13.592 | Lipid | Severe | *MAT2A* | Transcript | moderate | *RPGR* | Transcript | Mild |
| Unknown_mz_508.37686_+_RT_14.624 | Lipid | Severe | *MB* | Transcript | moderate | *S100A12* | Transcript | Mild |
| Unknown_mz_532.37421_+_RT_14.589 | Lipid | Severe | *MMP9* | Transcript | moderate | *S100A8* | Transcript | Mild |
| Unknown_mz_535.4339_+_RT_22.661 | Lipid | Severe | *NBN* | Transcript | moderate | *SERPINA1* | Transcript | Mild |
| Unknown_mz_552.40222_+_RT_14.023 | Lipid | Severe | *NR1H2* | Transcript | moderate | *SH2D3C* | Transcript | Mild |
| Unknown_mz_561.44934_+_RT_22.704 | Lipid | Severe | *NRL* | Transcript | moderate | *SIK1* | Transcript | Mild |
| Unknown_mz_568.33972_+_RT_11.902 | Lipid | Severe | *OSBP* | Transcript | moderate | *TGFB1* | Transcript | Mild |
| Unknown_mz_570.50861_+_RT_23.416 | Lipid | Severe | *PCYT1A* | Transcript | moderate | *TRIM21* | Transcript | Mild |
| Unknown_mz_614.3454_-_RT_12.087 | Lipid | Severe | *PECAM1* | Transcript | moderate | *VDR* | Transcript | Mild |
| Unknown_mz_616.36139_-_RT_12.605 | Lipid | Severe | *PLEK* | Transcript | moderate | *CCL2* | Protein | Mild |
| Unknown_mz_621.30347_-_RT_11.079 | Lipid | Severe | *PRF1* | Transcript | moderate | *IL13* | Protein | Mild |
| Unknown_mz_656.58331_+_RT_25.394 | Lipid | Severe | *PTGDR* | Transcript | moderate | 1-(1-enyl-palmitoyl)-2-linoleoyl-GPE (P-16:0/18:2)* | Metabolite | Mild |
| Unknown_mz_667.53271_-_RT_14.302 | Lipid | Severe | *PTPA* | Transcript | moderate | 1-(1-enyl-stearoyl)-2-docosapentaenoyl-GPE (P-18:0/22:5n3)* | Metabolite | Mild |
| Unknown_mz_684.61389_+_RT_26.119 | Lipid | Severe | *RAB1A* | Transcript | moderate | 1-(1-enyl-stearoyl)-2-linoleoyl-GPC (P-18:0/18:2)* | Metabolite | Mild |
| Unknown_mz_708.54657_+_RT_14.3 | Lipid | Severe | *RB1* | Transcript | moderate | 1-adrenoyl-GPC (22:4)* | Metabolite | Mild |
| Unknown_mz_708.62976_+_RT_27.454 | Lipid | Severe | *RIT1* | Transcript | moderate | 1-margaroyl-GPC (17:0) | Metabolite | Mild |
| Unknown_mz_712.6452_+_RT_25.192 | Lipid | Severe | *RPS20* | Transcript | moderate | 1-palmitoyl-2-stearoyl-GPC (16:0/18:0) | Metabolite | Mild |
| Unknown_mz_715.53271_-_RT_13.482 | Lipid | Severe | *SLA* | Transcript | moderate | 1-palmityl-2-linoleoyl-GPC (O-16:0/18:2)* | Metabolite | Mild |
| Unknown_mz_719.56488_-_RT_14.751 | Lipid | Severe | *SLC11A1* | Transcript | moderate | 1-palmityl-GPC (O-16:0) | Metabolite | Mild |
| Unknown_mz_724.30872_-_RT_23.18 | Lipid | Severe | *SOS1* | Transcript | moderate | 1-pentadecanoyl-GPC (15:0)* | Metabolite | Mild |
| Unknown_mz_738.65997_+_RT_26.549 | Lipid | Severe | *SREBF2* | Transcript | moderate | 1-stearyl-GPC (O-18:0)* | Metabolite | Mild |
| Unknown_mz_738.66125_+_RT_27.291 | Lipid | Severe | *SRF* | Transcript | moderate | 21-hydroxypregnenolone disulfate | Metabolite | Mild |
| Unknown_mz_738.6615_+_RT_25.394 | Lipid | Severe | *STAT5A* | Transcript | moderate | 6-bromotryptophan | Metabolite | Mild |
| Unknown_mz_739.58575_+_RT_26.834 | Lipid | Severe | *SYK* | Transcript | moderate | arachidonoylcholine | Metabolite | Mild |
| Unknown_mz_741.60077_+_RT_27.532 | Lipid | Severe | *TLR3* | Transcript | moderate | behenoyl sphingomyelin (d18:1/22:0)* | Metabolite | Mild |

We identified chemokines (*CXCL2, CXCL3, CXCR1, CXCR2, CXCR3, CXCR6),* cytokines (e.g., *TNF, TNFRSF1A, TNFSF10)* and other transcripts and proteins (e.g., *ATP6AP2*) that promotes the cytokine storm to discriminate the severe disease state from the other states. Concordantly, several studies have reported elevated levels of these features in COVID-19 patients, particularly those with severe disease (48–50). Also, some of the biosignatures including but not limited to *TNF, IL-10*, have been verified using multiplex biosensor techniques to be associated with COVID-19, and as such these biosignatures could serve as markers to monitor disease development (43, 51).

The results further revealed lysophosphatidylcholine (LysoPC), diacylglycerol (DG), and triglycerides (TG) to discriminate severe disease states. Several studies have suggested the possible differential association of these lipids with COVID-19 pathogenesis and disease severity (52–55).

Also, we identified metabolites such as kynurenate (56), gluconate (57), and sphingomyelin (58), which have been reported to play a role in the cytokine storm and immune response, to discriminate the moderate disease state from the other states.

Comparing the distribution of biosignatures discriminating disease states (**Figure 6A and Figure 6B**) further supports the idea that transcripts, metabolites, lipids, and proteins collectively influence disease progression beyond the mild state. Also, we identified features (e.g, *SLC14A1, Adipoylcarnitine*) for which no direct roles in influencing disease severity have previously been reported: further research may provide important insights into their roles in COVID-19 pathogenesis and whether they might be useful targets for therapeutic intervention. Overall, these findings suggest that the discriminatory features play a significant role in the immune response to COVID-19 and that targeting them and/or their associated signalling pathways may be a potential therapeutic approach (48).

**Figure 6.**
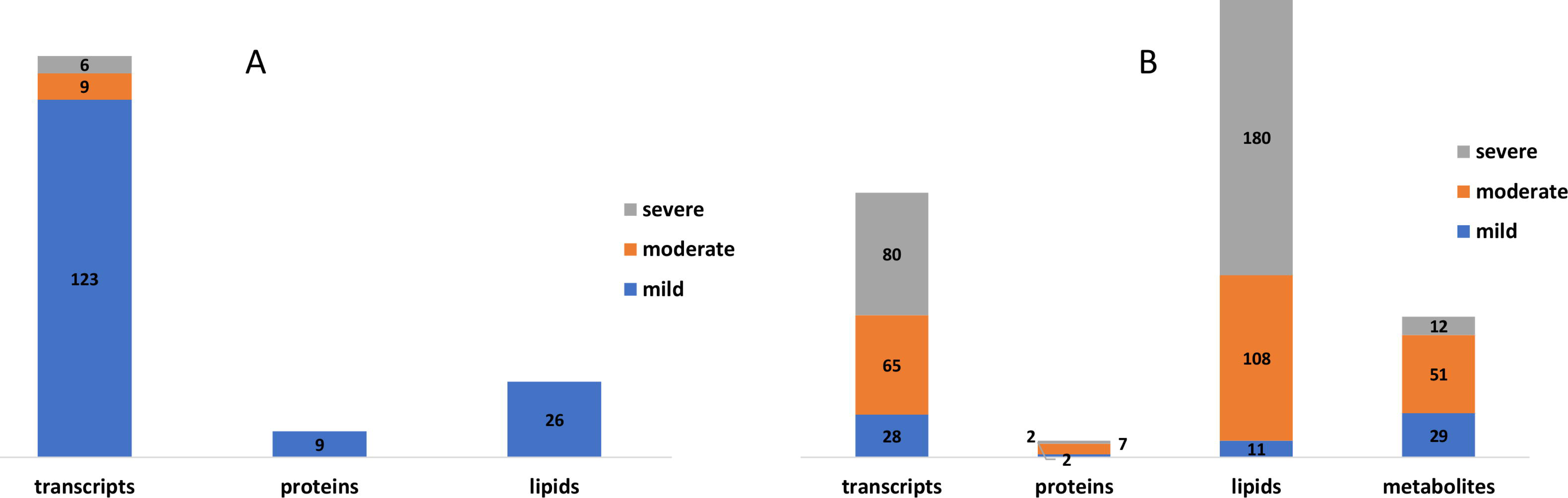
**(A)** Frequency of biosignatures based on omics feature type that discriminate COVID-19 disease states according to the analysis performed using the data-driven approach **(B)** Frequency of biosignatures based on feature type that discriminate COVID-19 disease states according to the analysis performed using the hypothesis-driven approach

### Enrichment analysis reveals enriched processes and pathways

#### Enrichment analysis of biosignatures that discriminate disease states based on data-driven seeds

From the discriminating features identified using the data-driven approach, we performed enrichment analysis based on the disease states they are differentially associated with (**Supplementary File 3**). The biological processes associated with proteins that discriminate the mild disease state is given in **Supplementary File 3 Table 1**. With a focus on the top 25 enriched biological processes and pathways, proteins were involved with chemotaxis (neutrophil, granulocyte, eosinophil, monocyte), regulating cytokine responses, and cell migration. These findings align with evidence of the role of chemotaxis in the initial response of detecting and destroying infected cells by following a chemical gradient of cytokines, chemokines, and other signaling molecules that are released by these cells (59). Further, the proteins were enriched in chemokine-mediated and interleukin-mediated pathways (**Supplementary File 3 Table 2**). For instance, regulation of the complement cascade pathway is important for controlling the immune response and preventing tissue damage. Several regulators of the complement system, including complement factor H, complement factor I, and CD59, are downregulated by SARS-CoV-2, which may contribute to complement dysregulation during infection (60, 61).

Transcripts discriminating the moderate disease state were enriched in those involved in cellular responses to organic substances and the mitogen-activated protein kinase (MAPK) cascade (**Supplementary File 3 Table 5**) but were especially enriched in transcripts involved in receptor-mediated signalling pathways (**Supplementary File 3 Table 6**). The MAPK pathways play a crucial role in regulating a variety of cellular responses, including cell proliferation, differentiation apoptosis, and immune response to COVID-19 (62).

The severe disease state discriminating transcripts were enriched in those involved in regulating ion transmembrane transporter activity, cell differentiation (dendritic, myeloid leukocyte), apoptotic processes, and mitochondrion organization (**Supplementary File 3 Table 7**) and enriched in signaling-related and regulatory-related pathways (**Supplementary File 3 Table 8**). The identified processes further align with the role of cell differentiation in COVID-19. For instance, dendritic cells act as sentinels to detect and respond to viral infections. They play a critical role in presenting viral antigens to T-cells, which, in turn, activate the immune response. During SARS-CoV-2 infections, dendritic cells can become infected which can lead to impaired antigen presentation and reduced activation of T-cells (63). Also, myeloid leukocytes, including monocytes and macrophages, are important in the early immune responses to COVID-19. These cells can phagocytose viral particles and present viral antigens to T-cells, activating the immune response (64). However, during severe infections, excessive activation of myeloid cells can lead to a cytokine storm (65), a dangerous immune response that can cause severe tissue damage and organ failure.

In addition to transcripts, there were also lipids that discriminated against the different disease states. We were, however, unable to perform enrichment analysis on these because all of the discriminating lipids are presently uncharacterized.

#### Enrichment analysis of biosignatures that discriminate disease states based on hypothesis-driven seeds

We repeated the enrichment analysis but with discriminatory features identified using hypothesis-driven seed selections (*IL6* and *IL6R*) (**Supplementary File 4 Tables 1-17**). The proteins and transcripts discriminating the mild, moderate, and severe disease states for example *IL13, CCL2*, *IL1A, SYK, IFNG, IL16, HMGB1, and TLR3* are involved in cytokine-, regulatory-mediated and apoptotic biological processes (**Supplementary File 4 Tables 1, 3, 7, 9, 14, and 17**) and were also enriched in pathways including interleukin-mediated signalling, cytokine-mediated signalling, and cellular responses to stimuli (**Supplementary File 4 Tables 2, 4, 8, 10, 15 and 17**). The fact that most of the biological processes and pathways are regulatory-, cytokine-, and cellular response-related agrees with other studies on disease severity. In the absence of appropriate regulatory T-cell activity to restrain the immune response to SARS-CoV-2 infections, the over-production of cytokines can ensue leading to a counter-productive cytokine storm (66, 67). Also, apoptotic biological processes are crucial in preventing severe disease by facilitating the death of infected cells so as to contain both the sizes of infection foci and the immune responses to the infected cells within these foci (67, 68).

The summary of metabolite pathways (**Supplementary File 4 Table 5**) linked to the mild disease states revealed metabolic processes that may play important roles in the pathophysiology of COVID-19 (69–71). For instance, sphingolipids are important components of cell membranes and are involved in a variety of cellular processes, including inflammation and apoptosis and the metabolism of these lipids has been implicated in the pathogenesis of viral infections, including COVID-19 (69, 72). Also, the dysregulation of arginine biosynthesis and lysine degradation may play a role in the pathogenesis of COVID-19 by modulating the immune response because arginine and lysine are essential amino acids that are involved in many biological processes including immune function and protein synthesis (71). We identified Sphingolipid metabolism processes to be common across all disease states. It has been suggested that the virus hijacks sphingolipid metabolism and dysregulates the metabolism activities to promote its replication and to evade the host immune response, aligning with the involvement of these processes in the pathogenesis of severe disease patients (**Supplementary File 4 Tables 5, 6, and 12**) (72). We also identified other pathways linked with metabolic pathways discriminating moderate and severe disease states including, but not limited to, phenylalanine, tyrosine, and tryptophan biosynthesis pathways, and the pentose phosphate pathway.

Given, that we could not perform enrichment analysis for uncharacterized lipids discriminating mild disease states, the summary of lipid pathways (**Supplementary File 4 Table 11 and 13**) linked with moderate and severe disease states revealed processes that may play important roles in the pathophysiology of COVID-19. Autophagy is involved in several other biological processes, including antigen presentation, cell death, and immune regulation to maintain or restore homeostasis. Dysregulation of these processes has been implicated in the pathogenesis of various diseases including COVID-19 and has even been presented as a target for therapeutics (73, 74). Arachidonic acid metabolism is the pathway responsible for the production of various bioactive lipids, including prostaglandins, leukotrienes, and thromboxanes. Dysregulation of arachidonic acid metabolism has been implicated in the pathogenesis of numerous diseases and syndromes, including inflammation, cancer, and cardiovascular disease (71).

Lipids are central components of cell membranes, such that dysregulation of pathways such as glycosylphosphatidylinositol (GPI) anchor biosynthesis and glycerophospholipid metabolism occurred in all disease states (**Supplementary File 4 Tables 11 and 13**). For instance, Glycosylphosphatidylinositol (GPI)-anchor biosynthesis - GPI-anchor biosynthesis is the process by which GPI-anchored proteins are synthesized and inserted into the plasma membrane. GPI-anchored proteins play critical roles in cell signaling, immune function, and development. Glycerophospholipid metabolism is the metabolic pathway responsible for the synthesis and degradation of glycerophospholipids, which are also essential components of cell membranes. Abnormalities in glycerophospholipid metabolism have been previously implicated in the pathogenesis of several diseases including COVID-19 (75, 76).

#### Validating the integrative network-based multi-omics-driven data approach and replicating results from independent data

We performed random walk network analysis on disease-state omics-specific graphs and specifically investigated the behaviour of a new multi-layer graph generated from different datasets from the perspective of network statistical parameters (**Supplementary data**). We retrieved published metabolomics, transcriptomics, and proteomics features reported to be associated with the various COVID-19 disease states. Specifically, transcriptomics features specific to mild, moderate, and severe disease states were retrieved from Alqutami et al., (12). Proteomics features specific to the moderate disease state were retrieved from Zhong et al., (14). Features specific to the mild and severe disease states were retrieved from Patel et al., (21) and Suhre et al., (20). We generated protein-protein interaction networks using GeneMANIA. Metabolites differentially associated with mild, moderate, and severe COVID-19 disease states were retrieved from Jia et al., (17). For each disease state, by using the metabolite KEGG IDs as inputs we constructed a metabolite-metabolite interactome using MetaboAnalyst 5.0 (77), which is a knowledge-driven multi-omics integration platform (https://www.metaboanalyst.ca/). We further constructed the metabolite-protein interactome for each disease state from MetaboAnalyst 5.0 by using the metabolite KEGG IDs and gene IDs of the features that were differentially associated with the different disease states. We also included the lipid interactome generated from Overmyer et al., (25) datasets. We repeated the random walk analysis using the data-driven and hypothesis-driven seeds; except that, since, hydroxyoctanoate was not a feature in the generated networks, it was excluded from the data-driven seeds. We also performed a statistical network analysis of the multi-layered graphs generated. We observed that network heterogeneity and characteristic path length metrics correlated with disease severity (**Table 4**). In accordance with the analyses that we previously performed on our multi-layered graphs, we observed that network density decreased with disease severity. The statistical analysis also supported the observation during our multi-layered network analyses that crosstalk between features across multiple omics layers (layers containing different feature types) relates to disease severity and could be a distinctive factor underlying the heterogeneity in disease severity among patients.

## Discussion

Different single omics (11–21) and multi-omics (7, 22-28) studies have been conducted to provide insights into disease severity. However, computational network-based integrative analysis that considers different omics profiles from multiple studies with existing biological knowledgebases to explore proteomics, transcriptomics, metabolomics, and lipidomics biosignatures and their connections across different disease phases, to help explain clinical heterogeneity are limited.

Here we hypothesized that (i) investigating biosignatures across COVID-19 disease phases would provide insights into the observed clinical heterogeneity and facilitate an understanding of factors associated with disease severity, and (ii) associations between the biosignatures within a biological network would permit the prioritization of those biosignatures that discriminate the disease states, which may, in turn, provide insights into drug research. We performed an integrative multi-omics network analysis by using proteomics, transcriptomics, metabolomics, and lipidomics data from Overmyer et al. (25), and Su et al., (24). We demonstrate an approach for harmonizing the clinical severity of COVID-19 patients from independent studies leveraging the WOS and patient clinical metadata. We believe that this approach forms the basis of classifying COVID-19 patients from independent multi-omics studies and allows for the grouping of omics experimental data into disease states to perform computational network-based integrative multi-source multi-omics analysis.

From the random walk network analysis on the disease-state omics-specific graphs, we notice some unique patterns in the cross-layer interactions for mild, moderate, and severe disease states. Specifically, random walk analysis using the hypothesis-driven seeds resulted in networks with both a greater variety of features (particularly metabolites and lipids) and more interactions between different feature types, than was achieved in networks generated using data-driven seeds. This observation might be partly attributable to the fact that *IL-6* and *IL-6R* (the two hypothesis-driven seeds that were used) have a profound role during the anti-inflammatory response to COVID-19. Compared to interactions established by *IL-6R* as a seed, we observed that *IL-6* as a seed established a network with more interactions between proteins, transcripts, and metabolites related to cell function and immune responses (accessible at http://cytoscape.h3africa.org). *IL-6R* as a seed yielded a network that primarily captured protein and transcript interactions. Notable among the metabolites interacting with *IL-6* is taurine, an amino sulfonic acid involved in the regulation of oxidative stress, which is known to play an important role in COVID-19. Taurine levels have been reported to decrease in COVID-19 patients which potentially modulates disease progression via its antiviral, antioxidant, anti-inflammatory, and vascular-related effects (78). Other metabolites interacting with *IL-6* included serotonin, a neurotransmitter, known to play important roles in the immune system and in regulating inflammatory responses (79).

From the data-driven seeds (**Table 1**), we observed that *STAT1* established connections with hubs *IRF1*, and *CCL4* in mild, and hub *IRF1* in both moderate and severe disease states. *SOD2* connects with hub *IRF1* in the mild, moderate, and severe disease state networks. Also, 3-hydroxyoctanoate interacts with 3-hydroxyexanoate and 3-hydroxydecanoate in all disease states. The random walk analysis provided not only an insight into network connectivity but also the results from the network statistical analysis (**Table 4**) were consistent and overlapped. Of note, the outcome from the random walk is based on the choice of seeds (**Table 1**) used for the random walk analysis. The analysis generated a multi-layered graph for each disease state based on the exploration of the seeds across the disease-state omics-specific graphs.

Despite the heterogeneity of COVID-19 disease outcomes, the individual mild, moderate, and severe disease states seem to have characteristic degrees to which transcript, protein, metabolite, and lipid features associatively interact both with themselves and with one another. Upon evaluating the multi-layered graphs for the various disease states, we identified several associative interactions that were present irrespective of disease state and a number that seemed to be specific for particular disease states (**Supplementary File 2**). These associations should be further analyzed to better understand the causal effects.

In general, we observed that transcript-transcript interactions were the most commonly detected across all the disease states whereas metabolite-metabolite and lipid-lipid interactions were least commonly detected. This observation could partly be attributed to the fact that between 4 and 16 times more individual transcript features are present in the transcriptomics experimental datasets than are present in the lipidome and metabolome datasets respectively. However, irrespective of these differences, we observed that major distinctions among disease states are a result of cross-layer interactions: with protein-metabolite interactions being particularly notable. Specifically, we observed an overall increase in cross-layer interaction with disease severity using both data-driven and hypothesis-driven seeds for network exploration. We tested network statistics to confirm the network behaviour across disease states in both the original datasets and the validating datasets and found that cross-layer interactions within networks could be a distinctive feature of severe COVID-19.

From the evaluation of interactions associated with one disease state, we identified biosignatures including but not limited to *MMP9, CCL5, S100A8, TNFRSF1A*, Plasmenyl-PE P-18:0_18:2_RT_22.986, cystathionine, 6-bromotryptophan, and PE 18:1_18:1_RT_20.883 that discriminate the various disease states (**Tables 5 and 6**). These biosignatures are differentially associated with the disease states (24, 25). Further, from the enrichment analysis of these discriminatory biosignatures (**Supplementary Files 3 and 4**), we notice cytokine-, regulatory-mediated and cellular responses to infection processes were apparent among disease-state discriminating transcripts and proteins identified from both the data-driven and the hypothesis-driven approach. This gives us a general overview that, despite the heterogeneity of COVID-19 disease outcomes, the biological processes and pathways underlying the disease could be related, but with varying expression levels of the biosignatures involved. As expected, we identified the metabolic processes related to the disease-discriminatory metabolites. However, other metabolite-related pathways relating to the degradation and/or synthesis of essential amino acids were noticeable among moderate and severe disease state discriminating metabolites (**Supplementary File 4 Tables 5, 6, and 12**). Knowing that these essential amino acid processes contribute to protein synthesis, therefore, suggests that the disturbance of protein synthesis could contribute to the severity of the disease.

We only used lipidomics experimental data from a single study and it is likely therefore that this part of our analysis was proportionally underpowered relative to that involving transcripts, proteins, and metabolites. Furthermore, most of the lipids used for the analysis were uncharacterized and we were therefore unable to perform enrichment analyses on them. In the multi-layered networks we created, we also did not discover any protein-lipid edge types. This may be attributable in part to the protein-lipid bipartite data and the seed exploration during random walk analysis. Furthermore, future investigations could consider incorporating not only characterized (annotated) lipids but also additional data types such as epigenomics, microbiomics, and immunomics. Moreover, from the analysis, we observed more transcript-transcript interactions as compared to other omics features. This observation is at least partly attributable both to the unevenness in the number of features measured in the different omics experiments (with the transcriptomics experiment examining between 5 and 100 times more features than other types of omics experiments) and the fact that more research efforts have been focused on transcriptomics analyses than on those of other omics types. Despite these limitations, our study has some obvious strengths and the results presented revealed biosignatures and their interactions related to disease severity. Having demonstrated that cross-layer interactions could be a distinctive feature, if not a hallmark, of severe COVID-19, it warrants deeper investigation into the potential causal relationships that these cross-layer interactions, have with disease progression: relationships that might illuminate ways to prevent and/or reverse this progression.

## Conclusion

In this work, we delved into the identification and characterization of biosignatures and their specific molecular features that underly various phases of COVID-19 disease, by using an integrative and network-based approach to analyze multi-omics data. We emphasized the critical importance of integrating multi-omics data, to elucidate the molecular dynamics responsible for the wide-ranging clinical presentations of COVID-19. This integration considers both prior knowledgebases and multi-omics data from independent studies.

The major contributions of this work are: (1) development of a new method to harmonize patient disease severity metrics by leveraging WOS in conjunction with patient metadata; (2) compilation of a comprehensive COVID-19 knowledge graph by assembling (interactome) data from various curated sources; (3) construction of disease-state omics-specific graphs by integrating curated proteomics, transcriptomics, metabolomics, and lipidomics datasets from two independent studies; and (4) identification of biosignatures and their associated interactions that are shared or unique to mild, moderate, and severe COVID-19 disease states.

Our study not only pinpoints biosignatures that distinguish between disease states, but also demonstrates a correlation between the severity disease states of COVID-19 and cross-layer interactions of proteins, transcripts, metabolites, and lipids.

We are confident that the presented multi-omics data harmonization and network-based analysis approach can also be applied to other diseases.

## Methods

### Study design and procedures

The approach for this study (**Figure 1**) consists of five main steps including: (1) data curation and pre-processing; (2) disease severity harmonization; (3) construction of disease-state omics-specific graphs; (4) multi-layer network-based random walk analysis; and (5) enrichment analysis.

### Data sources

#### Multi-omics experimental data

Two independent multi-omics experimental datasets were used in this study; quantified blood plasma transcript, protein, and metabolite count data from Su et al., (24) and transcript, protein, metabolite, and lipid count data from Overmyer et al., (25). For Su et al., (24) the study samples consisted of 139 COVID-19 patients (60 males and 79 females) and 258 healthy controls (**Supplementary data Table 1**). These enrolled COVID-19 patients had an age range from 18 to 89 years (median = 58). The Overmyer et al., (25) study samples were collected from 128 adult patients of which 102 (64 males and 38 females) were COVID-19-positive and 26 (13 males and 13 females) were negative (**Supplementary data Table 1**). There were no significant differences between the average ages of males and females in either the COVID-19 positive group –(61.3 years for females and 63.1 years for males; (p-value = 0.56; calculated using t-test) or the COVID-19 negative group (59.5 years for females and 67.0 years for males; p-value = 0.25).

#### Protein-protein interactome

1832 names of human genes associated with COVID-19 were retrieved from the DisGeNET database version 5 (80). GeneMANIA (81) was used to generate an interactome for 1692 of these genes (140 were unrecognized by the GeneMANIA database). Interactions between genes were based on co-expression, physical interaction, co-localization, shared protein domains, genetic interaction, or predictions from manual curation.

#### Metabolite-metabolite interactome

A co-expression network was constructed from the metabolomics measurement data. Highly correlated pairwise interaction scores >=0.7 were used to select components of the metabolite-metabolite interactome that was used for downstream analysis.

#### Lipid-lipid interactome

A co-expression network was constructed from the lipidomics measurement data. Highly correlated pairwise interaction scores >=0.7 were used to select components of the lipid-lipid interactome that was used for downstream analysis.

#### COVID-19 knowledge graph

We further included a multi-modal, knowledge model of COVID-19 pathophysiology (COVID-19 Knowledge Graph version 0.0.2, https://github.com/covid19kg/covid19kg) (82). The graph incorporates nodes, covering 10 entity types (e.g. proteins, genes, chemicals, and biological processes) and relationships between the nodes, and we considered only protein, genes, transcripts, lipids, and metabolites node types and their interactions for downstream analysis.

#### Cross-layer interactome

We retrieved protein-transcript, metabolite-protein, and lipid-protein associations from Su et al (24), and Overymyer et al (25), and used these to construct a bipartite graph for network analysis.

### Harmonizing the clinical severity of patients

Patient sample metadata from both the Su et al., (24), and Overmyer et al., (25) sources were used for disease severity harmonization. For Su et al., (24) metadata, the disease state linked to the samples was described using the World Health Organization (WHO) Ordinal Scale (WOS) based on specific categories and characteristics including (1) uninfected – no evidence of infection; (2) infected but ambulatory with no limitation of activities; (3) infected with limitation of activities but still ambulatory; (4) hospitalized with no or mild oxygen therapy; (5) hospitalized with oxygen administered by mask or nasal prongs; (6) hospitalized with non-invasive ventilation or high-flow oxygen; (7) hospitalized with intubation and mechanical ventilation; (8) hospitalized, with intubation and mechanical ventilation together with additional organ support; and (9) death.

In the Overmyer et al., metadata (25), disease severity was quantified using hospital-free days at day 45 (HFD-45) scores: a composite outcome variable that accounts for the length of hospital stay. The utility of the HFD-45 score is derived from the fact that severe COVID-19 patients are those that are admitted to the hospital the longest as they require ventilatory support, while those with the most extreme cases die during hospitalization (25). The variable assigns a zero value (0-free days) to patients with severe disease who remain admitted longer than 45 days or die due to respiratory deterioration while admitted, and higher values of HFD-45 to patients with shorter hospitalizations and milder disease severity.

To harmonize the clinical severity of patients, we used the WOS as the reference for classifying disease severity into three disease states, such that: (1) mild disease state represents COVID-19 patients with WOS 1-2, (2) moderate disease state represents COVID-19 patients with WOS 3-4, and (3) severe disease state represents COVID-19 patients with WOS 5-9.

The Overmyer et al. (25), sample metadata included the following variables: (1) ICU Status (an indicator variable of the patient’s ICU status), (2) HFD-45, (3) Acute Physiologic Assessment and Chronic Health Evaluation II (APACHE II) Score (an indicator variable ranging from 0 (best health) to 71 (worst health) based on physiologic variables, age, and health conditions), and (4) Mechanical Ventilation Status (MVS) (an indicator variable describing the patient’s mechanical ventilation status). There was a correlation between the HFD-45 and ICU status with APACHE II and MVS. These variables could feasibly be mapped onto the WOS scale, knowing that the WOS scale is primarily based on respiratory status and oxygen/ventilation support. Accordingly, we leverage this metadata to map characteristics of the Overmyer et al. (25) study patients onto the WOS. Specifically, we assigned: mild disease state to COVID-19 patients with HFD-45 between 29-45 with no time spent in the ICU, moderate disease state to COVID-19 patients with HFD-45 between 29-45 who spent time in the ICU, or a HFD-45 between 21-28 regardless of time spent in the ICU, and severe disease state to COVID-19 patients with HFD-45 less than 20 regardless of time spent in the ICU.

### Data pre-processing

We conducted a two-step data pre-processing operation on the omics experimental data using a custom script. Outlier and missing values were removed, and data were normalized. Samples with more than 20% missing data in a certain data type were excluded. Similarly, biological features such as mRNA expression, with more than 20% of values missing across patients were dropped from the data. Z-score normalization was then applied such that each feature of the data (samples as columns and features as rows) had an average of 0 and a standard deviation of 1.

#### Feature mapping to unified identifiers

To ensure that the feature labels were unified, transcript and protein identifiers were mapped to gene-level IDs using the UniProt (https://www.uniprot.org/) and NCBI databases (https://www.ncbi.nlm.nih.gov/gene/). Metabolite and lipid name descriptors were maintained for all analyses other than functional analysis for which KEGG or PubChem IDs were used.

#### Building a unified knowledge graph

We assembled a unified knowledge graph by merging the protein-protein interactome, the metabolite-metabolite interactome, the lipid-lipid interactome, and the extracted data from the COVID-19 knowledge graph (82).

### Building a disease-state omics-specific graph

Protein-protein, transcript-transcript, lipid-lipid, and metabolite-metabolite co-expression networks for the various COVID-19 disease states (i.e, mild, moderate, severe) were constructed based on an integrated unified knowledge graph, and the pre-processed omics data using an R script. From the Su et al., omics data, we constructed three coexpression networks (protein-protein, transcript-transcript, and metabolite-metabolite) for each disease state, as well as constructed four coexpression networks (protein-protein, transcript-transcript, lipid-lipid, and metabolite-metabolite) for each disease state from the Overmyer data. This was achieved by evaluating the correlation between the expression of each linked feature pair (i.e., transcript-transcript, protein-protein, metabolite-metabolite, lipid-lipid feature pairs). Given an edge e(i, j) in the unified knowledge graph (G) which linked feature i and feature j, let x_i_ = (x_i1_, …, x_in_) and x_j_ = (x_j1_,.. ., x_jn_) be the vectors of values in all samples for feature i and j respectively. Pearson’s correlations were used to estimate the individual relationships between features for every pre-processed omics measurement using an R script (https://github.com/francis-agamah/Multi-source-multi-omics-network-analysis). The interaction between feature pairs is represented by the Pearson Correlation Coefficient (PCC) ranging between −1 (perfect negative correlation) and 1 (perfect positive correlation), with values of zero representing no correlation. We further rescaled the PCC score for each pairwise interaction by computing the absolute value to attain positive scores ranging between 0 and 1.

The coexpression networks constructed from each single omics datasets formed the baseline for the network integration. Specifically, the co-expression networks of the same omics type constructed from the two independent studies were integrated by merging the networks to construct four omics-specific graphs (one for each omics type) for each of the three disease states.

### Random walk network analysis

#### COVID-19 disease state graph exploration by a random walk with restart

We adapted multiXrank (83), a random walk with restart on a multilayer network algorithm to explore the disease-state omics-specific graphs. This algorithm was chosen because it enables random walk with restart on any kind of multilayer network generated from different data sources as compared to other methods that are limited in the combination and heterogeneity of networks that they can handle (29). We modified the configuration script for the algorithm to accept four disease-state omics-specific graphs for the purpose of our analysis.

For network exploration on each disease state, the disease-state omics-specific graphs, cross-layer interactome, and seed nodes were used as inputs for the algorithm. Outputs were multi-layered graphs that described the exploration of the seed nodes across the different disease-state omics-specific graphs and a list of features in each disease-state omics-specific graph ranked according to their proximity to seed nodes.

The parameter values for global restart probability (set to 0.7), and inter-layer jump probability in a given disease-state omics-specific graph (set to 0.5), were maintained. The probability to restart in a specific layer of a specific disease-state omics-specific graph was set to 1: a setting that meant the disease-state omics-specific graph was classifiable as a monoplex network.

The probability to restart in a specific disease-state omics-specific graph was set to 0. This meant that the random walker stayed within the network within which it began with a probability equal to 1.

To achieve homogeneous exploration, the probability to jump across different disease-state omics-specific graphs was set to 0.25 in consideration of the four disease-state network layers.

Briefly, the first step of the algorithm is to create adjacency matrices for the input graphs, followed by computing different transition probabilities of the random walk with restart on the graphs. The probabilities are estimated based on the concept that an imaginary particle starts a random walk from the seed node to other nodes in the network. These different transition probabilities describe the walks within a graph and the jumps between graphs. A higher probability score (close to 1) suggests a higher likelihood to walk or jump between graphs.

#### Identifying seed nodes for multi-layered network exploration

To select seed nodes for the analysis, we implemented two approaches; (1) a data-driven approach where we selected as a seed the highest-ranked feature from each disease-state omics-specific graph based on a computed integrated node centrality metric score, and (2) a hypothesis-driven approach where we selected seeds based on previously reported differential associations with different COVID-19 disease states. For the data-driven approach, the features were ranked by leveraging the node degree, closeness, betweenness, and eigenvector centrality metrics to compute an integrated score (**see Supplementary data equations 1 to 3**).

#### Ranking candidate multi-omics features for COVID-19 disease states

All the network nodes were scored and ranked by the algorithm according to their proximity to the seed nodes. The computed score was the geometric mean of the node’s proximity to the seeds.

### Enrichment Analysis

#### Metabolite Pathway

Metabolite pathway enrichment analysis was performed using the MetaboAnalyst 5.0 online Pathway Analysis tool (77) (accessed 26 January 2023). Metabolite names were entered as KEGG IDs, and when necessary, metabolite names were automatically adjusted to match the nomenclature recognized by MetaboAnalyst (e.g., *Hydroxypropanoate* as *hydroxypropanoic acid*). Using high-quality KEGG metabolic pathways as the backend knowledgebase, we used the hypergeometric test to examine the overrepresentation of metabolites predefined in the KEGG pathway present in the queried metabolites. This approach determined whether a particular group of compounds was represented more than expected by chance within the user-uploaded compound list. In the context of pathway analysis, we evaluated if compounds involved in a particular pathway were enriched compared to random hits.

#### Lipid Pathway

We conducted lipid pathway enrichment analysis using the LIPEA online Pathway Analysis tool (84) (accessed 26 January 2023). Lipid names were entered as a compound list and, when necessary, these names were adjusted to match the nomenclature recognized by LIPEA.

#### Gene Ontology Analysis

Protein/transcript enrichment analysis was performed using the online Enrichr Gene Ontology Resource (85) (accessed 26 January 2023). The resource provides gene-set libraries made of a set of related genes which are associated with a functional concept such as a biological pathway or process (85). Gene ID identifiers were used as input for the enrichment analyses. The resource computes Fisher’s exact test followed by a correction based on a mean rank and standard deviation from the expected rank for each term in each gene-set library (85).

## Declarations

### Ethics approval and consent to participate

Not applicable

### Consent for publication

The authors have consented for the work to be published.

### Availability of data and materials

#### Data and code

- The scripts and processed data used for this study is accessible at https://github.com/francis-agamah/Multi-source-multi-omics-network-analysis
- A containerized workflow with expanded readme file to allow easy replication of this study is accessible at the same github repository.
- All other data generate during this study are included in this publication and its supplementary information files.

#### Online resource

We have hosted interactive networks on the website (http://cytoscape.h3africa.org). The graphs hosted on the website describes interactions between proteins (green nodes), transcripts (grey nodes), lipids (pink nodes), metabolites (cyan nodes) and seeds (yellow nodes) across mild, moderate and severe COVID-19 states. The edges describe protein-protein interactions (green edges), lipid-lipid interactions (pink edges), transcript-transcript interactions (grey edges), metabolite-metabolite interactions (cyan edges), protein-metabolite (blue edges), transcript-metabolites (blue edges).

#### Competing interest

The authors declare that the research was conducted in the absence of any commercial or financial relationships that could be construed as a potential conflict of interest.

#### Funding

This work was partially funded by an LSH HealthHolland grant to the TWOC consortium, a large-scale infrastructure grant from the Dutch Organization of Scientific Research (NWO) to the Netherlands X-omics initiative (184.034.019), and a Horizon2020 research grant from the European Union to the EATRIS-Plus infrastructure project (grant agreement: No 871096).

#### Authors’ contributions

FEA, EC, and PH conceived the study. FEA conducted the data analysis and prepared the first draft. PH, TE, MS, DM, and, EC supervised the work. TE, MS, EC, DM, and PH contributed to the revision of the article. All authors contributed to the article and approved the submitted version.

## Acknowledgments

Computations were performed using cloud computing facilities provided by ilifu (https://www.ilifu.ac.za/, South Africa) and High-Performance Computing from CHPC (https://www.chpc.ac.za/).

## Notes

### Competing Interest Statement

The authors have declared no competing interest.

