## Supplementary data for "Network-based integrative multi-omics approach reveals biosignatures specific to COVID-19 disease phases"

^1^Computational Biology Division, Department of Integrative Biomedical Sciences, Institute of Infectious Disease and Molecular Medicine, Faculty of Health Sciences, University of Cape Town, Cape Town, South Africa. ^2^Medical BioSciences department, Radboud University Medical Center Nijmegen, The Netherlands. ^3^Department of Applied Sciences, Faculty of Health and Life Sciences, Northumbria University, Newcastle, Tyne and Wear, NE1 8ST, UK

*joint corresponding authors

Address correspondence to:

Peter A.C. 't Hoen, PhD

Medical BioSciences department

Radboud University Medical Center Nijmegen, The Netherlands

PO Box 9101, 6500 HB Nijmegen, The Netherlands

Emile R. Chimusa, MSc, PhD

Department of Applied Sciences,

Faculty of Health and Life Sciences,

Northumbria University, Newcastle,

Tyne and Wear, NE1 8ST, UKE

### **Materials and Methods**

**Table 1**: Description of multi-omics experimental data used for this study

| **Reference** | **Date (online)** | **Study title** | **Study size** | **Sample type** | **Longitudinal study?** | **Clinical data** | **Describing heterogenuos disease trajectory** | **OMICS technologies used** | **OMICS other remarks** | **OMICS sample size** |
| --- | --- | --- | --- | --- | --- | --- | --- | --- | --- | --- |
| **Su et al., Cell** | Oct 28, 2020 | Multi-Omics Resolves a Sharp Disease-State Shift between Mild and Moderate COVID-19 | 139 patients ; 258 healthy controls ; | blood plasma | YES : ONLY for patients: n = 2 ; First blood sample was collected shortly after the initial clinical diagnosis (t1), Second was collected a few days later (t2) | COVID-19 status ;  disease severity (WOS) ; age, sex, BMI, race/ethnicity ; more clinical metadata available in the supplement | WHO ordinal score The score was used to classify patients along their disease course | proteomics | 464 proteins measured | 135 patients with 259 samples, 124 healthy controls with sample |
|  |  |  |  | blood plasma |  |  |  | metabolomics | 1050 metabolites measured | 133 patients with 254 samples, 133 healthy controls with sample |
|  |  |  |  | blood PBMC |  |  |  | transcriptomics | whole transcriptomics ;  single-cell | 0.55M cells from 254 patient samples, 16 healthy control samples |
|  |  |  |  | blood PBMC |  |  |  | secretomics | cytokine panel ; single-cell | 2 different multiplex cytokine panels with 32 proteins each, one for Innate and one for Adaptive Immunity, with 101K cells from 50 patient samples, 7 healthy control samples |
|  |  |  |  | blood PBMC |  |  |  | surface markers | TotalSeq-C ; single-cell | 0.55M cells from 254 patient samples, 16 healthy control samples (with TotalSeq-C, i.e. done in the same run as the transcriptomics analysis) |
|  |  |  |  | blood PBMC |  |  |  | TCR sequencing | T (and B) cell receptor gene sequencing ; single-cell | 0.55M cells from 254 patient samples, 16 healthy control samples (done in the same run as the transcriptomics analysis) |
| **Overmyer et al., 2020, Cell** | 20-Jan-21 | Large-Scale Multi-omic Analysis of COVID-19 Severity | A cohort study involving 102 COVID-19 patients ; 26 non-COVID-19 patients | blood plasma | NO: Blood samples were collected from 128 adults admitted to Albany Medical Center at the time of enrollment | COVID-19 status ; disease severity (Hospital-free days at day 45 [HFD-45] and WHO ordinal score); clinical data detailing days admitted pre-enrollment, sex, age, ethnicity, disease severity indexes (charlson comorbidity index, acute physiologic assessment and chronic health evaluation (APACHE II) score, sequential organ failure assessment (SOFA) score, SAPPSII), biomarkers, hemogram, respiratory parameters and treatment are available in Table 1 | Hospital-free days at day 45 [HFD-45] and WHO ordinal score The score was used to classify patients along their disease course | metabolomics | discovery metabolomics targeted metabolomics; 110 meatabolites measured | 128 blood samples from 128 patients 13 plasma metabolites were significantly associated with COVID-19 status |
|  |  |  |  | blood plasma |  |  |  | lipidomics | Mass spectrometry-based assay -discovery lipidomics; 646 lipids measured | 168 plasma lipids were statistically associated with COVID-19 status |
|  |  |  |  | blood plasma |  |  |  | proteomics | Mass spectrometry-based assay - Shortgun proteomics; 517 proteins measured | 146 plasma proteins were significantly associated with COVID-19 status |
|  |  |  |  | leukocytes derived from paitient blood samples |  |  |  | transcriptomics | transcriptomes of leukocytes; 13263 transcripts measured | 2537 leucocyte transcripts were significantly associated with COVID-19 status |
|  |  |  |  |  |  |  |  |  | Mass spectrometry-based assay 2786 unidentified small molecules | 511 unidentified metabolites and lipids were significantly associated with COVID-19 diagnosis |

### **Materials and Methods**

### ***Identifying seed nodes for multi-layered network exploration***

#### The node degree, closeness, betweenness and eigenvector centrality metrics were computed using the igraph package in R. The integrated centrality score measures the difference among centralities for each node and describes the average distance.

Let $C_{i}^{j}$ represent the centrality score calculated for each node $V_{j}(j=1,\ldots,n_{v})$ in a disease state graph, $G_{i}$. The integrated node centrality score is calculated using **equation 1**.

$\theta=\frac{\sum_{i=1}^{r} D_{i}^{j}}{r}$*Equation 1*

Where $r$ is the number of centrality measures calculated

$D_{i}^{j}$ (**Equation 2**) is the distance between the centrality of nodes in each graph.

$D_{i}^{j}=\left| C_{i}^{j}-M^{j} \right|$*Equation 2*

Where $M^{j}$ is the average node centrality score as shown in **equation 3**

$M^{j}=\frac{\sum_{i=1}^{r} C_{i}^{j}}{r}$*Equation 3*

**Results**

#### **Harmonized clinical severity between patients' metadata**

**Table 2**: Summary of transcriptomics, proteomics, metabolomics and lipidomics samples per disease state after harmonization

| Disease state | Transcriptomics samples | Proteomics samples | Metabolomics samples | Lipidomics samples |
| --- | --- | --- | --- | --- |
| Severe | 73 | 94 | 95 | 44 |
| Moderate | 71 | 121 | 120 | 20 |
| Mild | 85 | 147 | 142 | 39 |

### **Characterizing multi-layered graphs**

In this section, we used statistical network measures to compare and characterize the generated multi-layered graphs. Specifically, we implemented network density, network heterogeneity, and characteristic path length statistical measures. Network density measures how sparse or dense a graph is according to the number of connections per node [1]. The closer the value is to 1, the denser the network. Biologically, a network of features of the same type (homogeneous network) turns to form more clusters, thus likely to have network density values closer to one compared to a heterogeneous network. This observation is based on the hypothesis that the features that cluster together share some biological functionality. The network heterogeneity quantifies the diversity of connections between nodes [2]. Biologically, the network heterogeneity provides insights into cellular heterogeneity and the level of graph connectivity (with other feature types) that can guide the selection of biomarkers. Interestingly, network heterogeneity provides an overview of disease classification in terms of the transition from one disease state to another [3]. The characteristic path length measures the average number of edges in the shortest paths, particularly between node pairs of the same type [4]. Biologically, this measure also provides insights to graph connectivity. In the case of an unconnected graph, the characteristic path length is infinite.
