## Supplementary figures and images for "Network-based integrative multi-omics approach reveals biosignatures specific to COVID-19 disease phases"

### Supplementary Figure 1

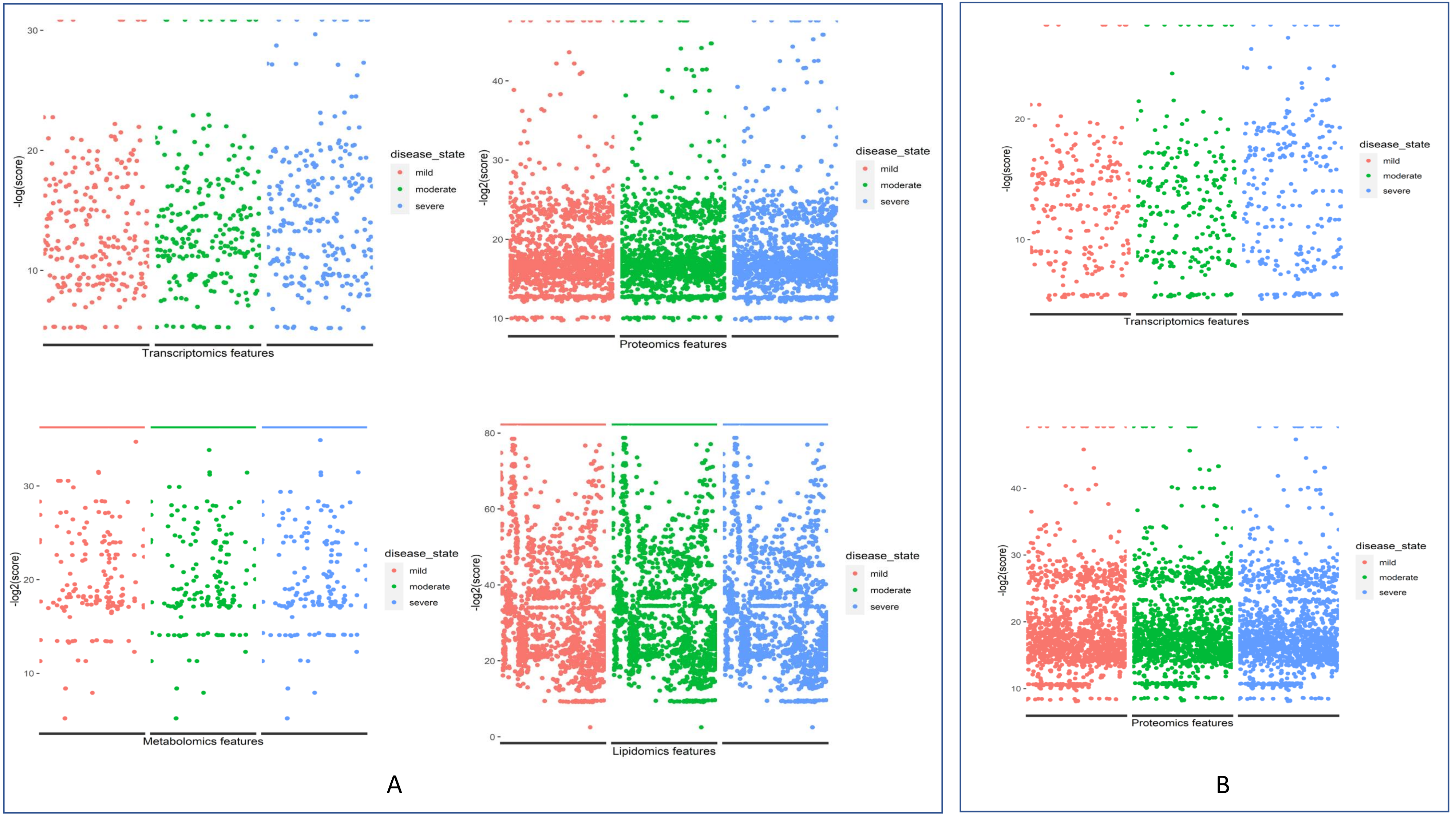
